# Inflammasome-mediated antagonism of type I interferon enhances *Rickettsia* pathogenesis

**DOI:** 10.1101/697573

**Authors:** Thomas P. Burke, Patrik Engström, Roberto A. Chavez, Joshua A. Fonbuena, Russell E. Vance, Matthew D. Welch

**Affiliations:** Department of Molecular and Cell Biology, University of California, Berkeley, Berkeley, CA, 94720 USA; Howard Hughes Medical Institute

**Keywords:** Inflammasome, type I interferon, IFN-γ, *Rickettsia parkeri*, IRF5, guanylate-binding proteins, GBP2, GBP5, caspase-11, cGAS

## Abstract

Inflammasomes and interferons constitute two critical arms of innate immunity. Most facultative bacterial pathogens that inhabit the host cell cytosol avoid activating inflammasomes and are often resistant to killing by type I interferon (IFN-I). We report that the human pathogen *Rickettsia parkeri,* an obligate intracellular pathogen that resides in the cytosol, is sensitive to IFN-I. The mechanism of IFN-I-dependent restriction requires the transcription factor IRF5, which upregulates anti-rickettsial factors including guanylate-binding proteins and iNOS. However, *R. parkeri* curtails cGAS-dependent IFN-I production by causing caspase-11-dependent pyroptosis. *In vivo*, inflammasome activation antagonizes IFN-I production, enhancing *R. parkeri* abundance in the spleen. Mice lacking either IFN-I or IFN-γ signaling are resistant to infection, but mice lacking both rapidly succumb, revealing that both interferons are required to control *R. parkeri*. This study illuminates how an obligate cytosolic pathogen exploits the intrinsic trade-off between cell death and cytokine production to escape killing by innate immunity.

**Highlights:**

- *Rickettsia* killed by GBPs activates caspase-11 and GSDMD, promoting pyroptosis
- *Rickettsia* exploits pyroptosis to avoid cGAS-dependent type I interferon
- IRF5, GBPs, and iNOS contribute to controlling *R. parkeri* infection
- *Ifnar*^-/-^*Ifngr*^-/-^ mice succumb to infection, uncovering a mouse model to study *R. parkeri*

## Introduction

The innate immune response to microbial pathogens depends on upregulation of antimicrobial factors, secretion of cytokines, and activation of host cell death pathways (Jorgensen and Miao, 2015; Meunier and Broz, 2016; Mitchell and Isberg, 2017; Randow et al., 2013). Innate immune responses have been characterized during infection with facultative intracellular bacterial pathogens such as *Listeria monocytogenes* and virulent *Francisella* species, as well as viruses, which inhabit the host cell cytosol (Brubaker et al., 2015; McNab et al., 2015; Wallet et al., 2016). However, less well understood are the innate immune responses to obligate intracellular bacterial pathogens. Spotted fever group (SFG) *Rickettsia* spp. are tick-borne pathogens that cause spotted fever diseases worldwide (Walker and Ismail, 2008). As obligate intracellular bacteria that replicate exclusively in the host cell cytosol, SFG *Rickettsia* spp. must continually interface with host innate immune sensors. Therefore, they presumably have evolved sophisticated mechanisms to avoid or even exploit innate immune responses.

Following infection and subsequent detection of pathogen associated molecular patterns (PAMPS) by host cell sensors (Takeuchi and Akira, 2010), cytokines including type I interferon (IFN-I) are upregulated. For example, the detection of cytosolic DNA by cyclic GMP-AMP synthase (cGAS) results in activation of the downstream adaptor stimulator of IFN genes (STING) and transcription factors including IFN regulatory factor 3 (IRF3) to upregulate IFN-I expression and secretion (Sun et al., 2013). Binding of IFN-I to the interferon-α/β receptor (IFNAR) then activates a signaling cascade that upregulates the expression of hundreds of interferon-stimulated genes (ISGs), mobilizing the cytosol to an antimicrobial state (MacMicking, 2012; Meunier and Broz, 2016).

Previous studies revealed that *L. monocytogenes* is resistant to the killing effects of IFN-I in macrophages (Reutterer et al., 2008; Woodward et al., 2010), and in fact, this pathogen actively secretes STING agonists that stimulate IFN-I production (Burdette et al., 2011; Woodward et al., 2010). IFN-I enhances *L. monocytogenes* as well as *Francisella novicida* pathogenicity *in vivo* (Auerbuch et al., 2004; Carrero, 2013; Henry et al., 2010; O’Connell et al., 2004; Storek et al., 2015), demonstrating that these bacterial pathogens benefit from IFN-I signaling. In contrast, IFN-I has a critical role in restricting viral replication, and therefore the effects of IFN-I signaling on pathogens that inhabit the cytosol are often regarded as anti-viral but not anti-bacterial (Boxx and Cheng, 2016; McNab et al., 2015; Stetson and Medzhitov, 2006).

Cytosolic PAMPs can also be recognized by inflammasomes, resulting in host cell death. For example, sensing of PAMPs such as microbial DNA by nucleotide-binding domain and leucine-rich repeat containing gene family (NLR) proteins causes their oligomerization and subsequent activation of the protease caspase-1 (Lamkanfi and Dixit, 2014; Strowig et al., 2012). Similarly, direct binding of bacterial lipopolysaccharide (LPS) to the non-canonical inflammasome caspase-11 causes its oligomerization and activation (Aachoui et al., 2013; Hagar et al., 2013; Kayagaki et al., 2013; Shi et al., 2014). Active caspases-1 and −11 cleave the pore-forming protein gasdermin D (GSDMD), promoting pyroptosis, a rapid lytic host cell death (Kayagaki et al., 2015; Shi et al., 2015). Pyroptosis curtails microbial replication and exposes microbes to the extracellular environment where they can be targeted by immune factors such as phagocytes and antibodies. To avoid inflammasome activation, *F. novicida* modifies its LPS, enabling it to avoid caspase-11 (Hagar et al., 2013; Wallet et al., 2016). Additionally, *L. monocytogenes* and *F. novicida* minimize bacteriolysis to prevent the release of DNA (Peng et al., 2011; Sauer et al., 2010), and *L. monocytogenes* downregulates expression of flagellin (an activator of the inflammasome) during infection (Shen and Higgins, 2006). These virulence strategies reveal that pathogens can benefit from circumventing inflammasome activation. Interestingly, inflammasome activation antagonizes IFN-I production (Banerjee et al., 2018; Corrales et al., 2016; Jabir et al., 2014; Liu et al., 2018), and IFN-I can antagonize inflammasome activation (Guarda et al., 2011; Inoue et al., 2012). This suggests that the immune system may flip a switch between cell-intrinsic anti-viral and anti-bacterial responses.

Despite these advances, the role for IFN-I, inflammasomes, and other innate immune responses during infection by obligate intracellular bacteria is not fully understood. IFN-I has modest anti-rickettsial effects in immortalized endothelial cells *in vitro* (Colonne et al., 2011; 2013; Turco and Winkler, 1990). In regards to the inflammasome, caspase-1 is required for the production of IL-1β during *R. australis* infection (Smalley et al., 2016). Many studies have also established a critical role for interferon-γ (IFN-γ) and nitric oxide as anti-rickettsial factors (Feng et al., 1994; Li et al., 1987; Turco and Winkler, 1993; Turco et al., 1998; Walker et al., 1997). However, the role for pyroptosis, IFN-I, and their relationships to IFN-γ during *Rickettsia* infection remain unknown. Since *Rickettsia* obligately reside within the host cytosol, we hypothesize that they have evolved sophisticated and distinctive measures to manipulate the inflammasome and IFN-I response to their benefit.

Here, we investigated the roles for host cell death, IFN-I signaling, and antimicrobial factors in controlling *R. parkeri* infection at the molecular, cellular, tissue, and organismal levels. We observed that, unlike facultative cytosolic bacterial pathogens, *R. parkeri* intracellular growth is restricted by IFN-I in macrophages *in vitro*. If *R. parkeri* is killed in the host cytosol by guanylate-binding proteins (GBPs), the inflammasome is activated, thus curtailing IFN-I production. Consistently, we discovered that *R. parkeri* antagonized IFN-I production *in vivo* via inflammasome activation, which has tissue-specific effects on controlling bacterial replication. We also found that both IFN-I and IFN-γ are critical for controlling *R. parkeri* infection in mice, and propose that mice lacking interferon signaling can serve as a robust animal model for investigating *Rickettsia* pathogenesis. Our data suggest an unprecedented strategy whereby a bacterial pathogen exploits the inherent antagonism between the inflammasome and IFN-I response to promote its pathogenesis.

## Results

### Inflammasome activation promotes *R. parkeri* growth in macrophages by antagonizing the IFN-I response

Although previous studies have investigated the roles of IFN-I and inflammasomes during infection by facultative intracellular pathogens, little is known for obligate intracellular pathogens including *R. parkeri*. Previous reports suggested that IFN-I causes a ∼50% reduction in *Rickettsia* spp. growth in immortalized endothelial cells (Colonne et al., 2011; 2013; Turco and Winkler, 1990). *In vivo*, both endothelial cells (Feng et al., 1993; Sahni and Rydkina, 2009) and other cell types including macrophages (Banajee et al., 2015; Feng et al., 1993; Osterloh et al., 2016; Papp et al., 2016; Walker et al., 1999) are targeted by *Rickettsia* spp. during infection. However, the effect of IFN-I on *Rickettsia* spp. replication in primary macrophages was unknown. We infected wild type (WT) primary bone marrow-derived macrophages (BMDMs) with *R. parkeri* or *L. monocytogenes*, treated them with recombinant IFN-β, and measured the impact on bacterial growth (by counting plaque-forming units (PFU) for *R. parkeri* or colony-forming units (CFU) for *L. monocytogenes*). IFN-β caused a dose-dependent restriction of *R. parkeri* growth (**Figure 1A**), whereas it had no effect on *L. monocytogenes* (**Figure 1B**). We further assessed whether each of these pathogens elicited an IFN-I response during infection by testing supernatants from infected BMDMs for their ability to stimulate expression of an IFN-responsive luciferase reporter. *R. parkeri* infection did not induce appreciable IFN-I expression, whereas *L. monocytogenes* elicited a robust response (**Figure 1C**). These data suggest that, although *R. parkeri* is sensitive to IFN-I, it elicits low amounts of IFN-I and therefore grows robustly in macrophages.

**Figure 1:**
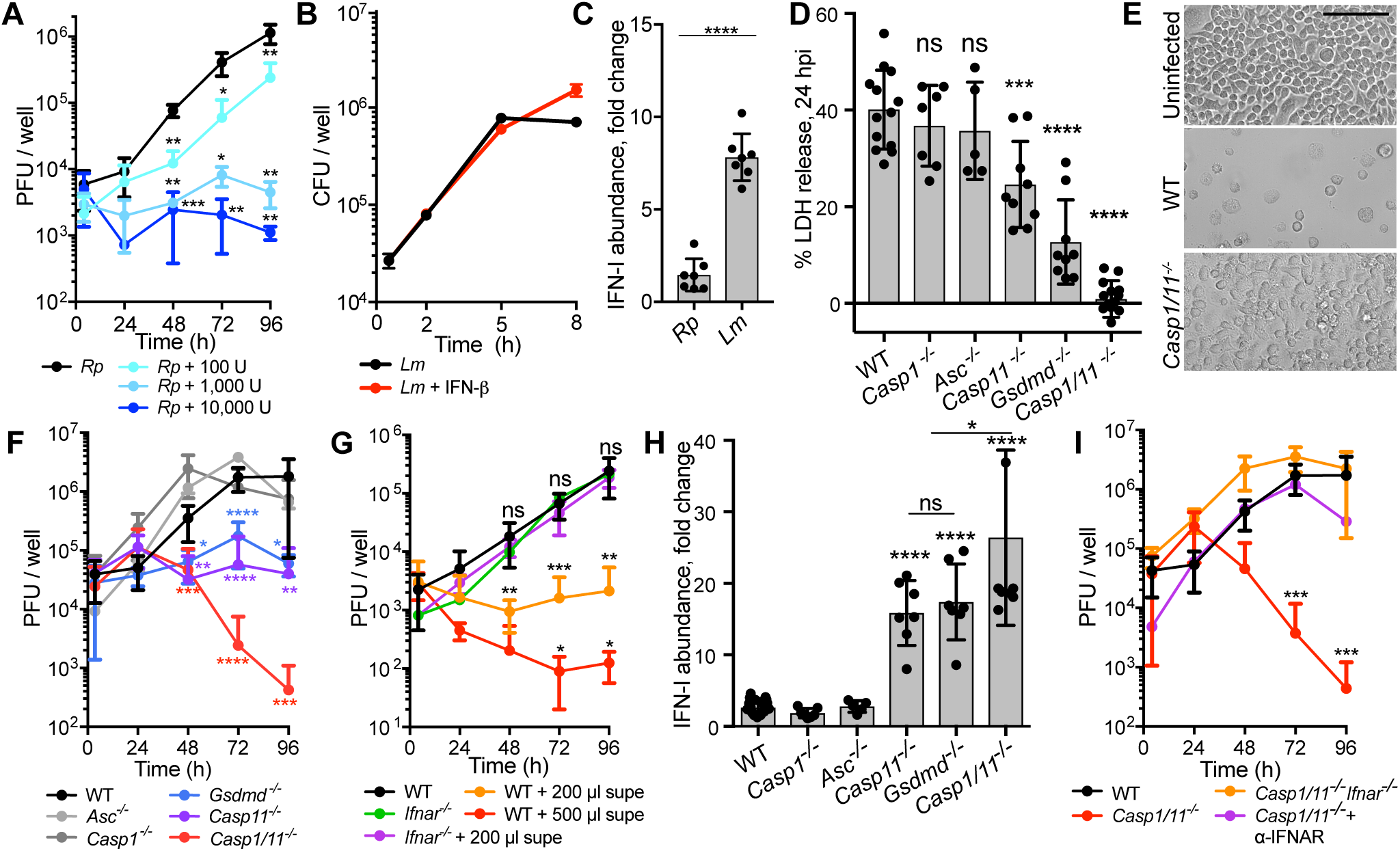
Inflammasome activation promotes *R. parkeri* pathogenesis by antagonizing the IFN-I response in macrophages. **A**) Measurement of *Rickettsia parkeri* (*Rp*) PFU in BMDMs over time. WT BMDMs were infected in a 24-well plate at an MOI of 0.2. MOI was calculated based on the ratio of the number of infectious bacteria, as determined by plaque assay, to the number of BMDMs. At each time point, BMDMs were lysed with water and lysates were plated over a confluent monolayer of Vero cells. Recombinant IFN-β (PBL, 12405-1) was added at 0 hpi. U signifies units of recombinant IFN-β. Statistical comparisons were made between each sample and untreated cells at each time point. Data are a compilation of at least two separate experiments and are expressed as means ± SEM. **B**) Measurement of *Listeria monocytogenes* (*Lm*) CFU in BMDMs. WT BMDMs were infected in 24-well plate at an MOI of 1. MOI was calculated based on the ratio of bacteria in liquid culture to BMDMs. 3,000 U of recombinant IFN-β (PBL, 12405-1) was added at 0 hpi. Data are the combination of at least two separate experiments and are expressed as means ± SEM. **C**) Measurement of IFN-I in supernatants of cells infected with *R. parkeri* or *L. monocytogenes*. WT BMDMs were infected with *R. parkeri* or *L. monocytogenes* at an MOI of 1 and supernatants were harvested at 8 hpi for *L. monocytogenes* and 24 hpi for *R. parkeri.* Supernatants were used to stimulate a luciferase-expressing cell line and relative light units (RLU) were measured and compared between each sample and uninfected cells. Statistical comparisons were made between each sample and infected WT cells. Data are the compilation of at least two separate experiments and are expressed as means ± SEM. **D**) Quantification of host cell death during *R. parkeri* infection of BMDMs. LDH release was measured at 24 hpi upon *R. parkeri* infection at an MOI of 1. Statistical comparisons were made between each sample and WT B6 cells. Data are the compilation of at least three separate experiments and are expressed as means ± SEM. **E**) Images of BMDMs infected with *R. parkeri*. Cells were infected at an MOI of 1 and images were captured at 96 hpi. The scale bar is 100 μm. **F**) Measurement of the number of infectious *R. parkeri* in BMDMs over time. BMDMs were infected with *R. parkeri* at an MOI of 1 and PFU were determined every 24 hpi. Statistical comparisons were made between each sample and WT B6 cells at each time point. Data are the compilation of at least three separate experiments and are expressed as means ± SEM. **G**) Measurement of the number of infectious *R. parkeri* in BMDMs over time. Cells were infected with *R. parkeri* at an MOI of 0.2 and PFU were measured over time. Statistical comparisons were made between each sample and WT B6 cells at each time point. Data are the compilation of at least three separate experiments and are expressed as means ± SEM. “Supe” indicates supernatant collected at 24 hpi from *Casp1^-/-^Casp11^-/-^* BMDMs infected with *R. parkeri* at an MOI of 1. **H**) Measurement of IFN-I in supernatants of cells infected with *R. parkeri*. Supernatants from infected cells were harvested at 24 hpi and used to stimulate a luciferase-expressing cell line and relative light units (RLU) were compared between each sample and uninfected cells. Statistical comparisons were made between each sample and infected WT cells. Data are the compilation of at least three separate experiments and are expressed as means ± SEM. **I**) Measurement of the number of infectious *R. parkeri* in BMDMs over time. Cells were infected at an MOI of 1 and PFU were measured over time. The α-IFNAR antibody was added at T=0 to a final concentration of 1 µg/ml. Statistical comparisons were made between each sample and WT B6 cells at each time point. Statistical analyses for data in panels A, B, C, F, G, and I were performed using a two-tailed Student’s T-test, where each sample at each time point was compared to the control. Statistical analyses in panels D and H were performed using an ANOVA with multiple comparisons post-hoc test. *p<0.05, **p<0.01, ***p<0.001, ****p<0.0001, ns=not significant.

We next investigated how *R. parkeri* escapes inducing IFN-I by examining the interaction between *R. parkeri* and the inflammasome, leveraging the ability to generate BMDMs from mice genetically deficient in inflammasome components. First, we focused on caspase-1, apoptosis-associated speck-like protein containing CARD (ASC), and the non-canonical inflammasome component caspase-11. WT and mutant BMDMs were infected with *R. parkeri* and bacterial-induced cell death, a readout of inflammasome-dependent pyroptosis, was monitored at 24 h post infection (hpi) using a lactate dehydrogenase (LDH) release assay. Infected *Casp1^-/-^* and *Asc^-/-^* mutant BMDMs exhibited ∼40% death, similar to WT cells (**Figure 1D**). In contrast, cell death was significantly reduced in *Casp11^-/-^* mutant BMDMs, and little cell death was observed in *Casp1^-/-^Casp11^-/-^*double mutant BMDMs (**Figure 1D**). Because caspases-1 and −11 activate pyroptosis by cleaving the pore-forming protein GSDMD, we next measured host cell death in *Gsdmd^-/-^*BMDMs. Compared to *Casp1^-/-^* or *Casp11^-/-^* single mutants, *Gsdmd^-/-^*BMDM cell death was reduced, but it was not abolished as in the *Casp1^-/-^Casp11^-/-^*double mutant cells (**Figure 1D**). These data reveal that the majority of *R. parkeri-* induced host cell death in macrophages depends on caspase-11 and GSDMD. To a lesser extent, *R. parkeri* activates caspase-1-dependent and GSDMD-independent cell death.

As infections proceeded to 96 h, we observed that WT BMDMs became rounded and non-adherent, whereas many *Casp1^-/-^Casp11^-/-^* double mutant cells remained adherent, similar to uninfected cells (**Figure 1E**). When we measured the number of infectious bacteria in these cells by counting PFU, we observed a reduction in the number of bacteria in *Casp1^-/-^Casp11^-/-^* BMDMs, whereas the number of bacteria increased over time in WT BMDMs (**Figure 1F**). There was also a reduction of infectious bacteria in *Casp11^-/-^*and *Gsdmd^-/-^* single mutant BMDMs, albeit less pronounced than for the *Casp1^-/-^ Casp11^-/-^* double mutants (**Figure 1F**). Thus, a subpopulation of *R. parkeri* appears to exploit activation of caspase-11 and GSDMD to promote replication of the remaining population.

Cytokines including IFN-I and TNF-α are produced by macrophages and can exert antimicrobial activity (Carrero, 2013; Weiss and Schaible, 2015). In addition, IFN-I is overproduced in cells lacking the inflammasome (Corrales et al., 2016; Liu et al., 2018). Hence, we hypothesized that *R. parkeri-* infected *Casp1^-/-^Casp11^-/-^*BMDMs were secreting increased levels of IFN-I that restricted *R. parkeri* growth. We therefore tested the bactericidal effects of adding supernatants from infected *Casp1^-/-^ Casp11^-/-^* cells to WT BMDMs that were infected with *R. parkeri.* We found that this supernatant restricted bacterial growth in a dose-dependent manner (**Figure 1G**). Furthermore, IFN-I secretion was increased at least 15-fold in infected *Casp1^-/-^Casp11^-/-^* double mutant and *Casp11^-/-^* and *Gsdmd^-/-^* single mutant cells (**Figure 1H**). To test whether IFN-I secreted from infected *Casp1^-/-^Casp11^-/-^*BMDMs was necessary for bacterial killing, we transferred the supernatant from infected *Casp1^-/-^Casp11^-/-^* to infected *Ifnar^-/-^* BMDMs, which lack the IFN-I receptor and hence are not responsive to IFN-I. The supernatant had no effect on bacteria growth in *Ifnar^-/-^* cells (**Figure 1G**). Supernatants from infected *Casp1^-/-^Casp11^-/-^* cells were also treated with an anti-IFNAR antibody to prevent IFN-I signaling, and this similarly restored bacterial growth (**Figure 1I**). To further test whether IFN-I was the inhibitory factor, we bred mice lacking *Ifnar*, *Casp1,* and *Casp11 (Casp1^-/-^Casp11^-/-^Ifnar*^-/-^), infected BMDMs from these mice, and observed that growth of *R. parkeri* was restored (**Figure 1I**). A similar series of experiments failed to show a role for TNF-α in killing *R. parkeri* (**Figure S1**). Together, these findings demonstrate that activation of the inflammasome allows *R. parkeri* to avoid killing by IFN-I.

### Pyroptosis masks cGAS-induced IFN-I

It remained unclear whether activation of host immune sensors was due to release of *R. parkeri* ligands such as LPS or DNA. We therefore sought to identify the host sensor required for the increased IFN-I production observed in infected *Casp1*^-/-^*Casp11*^-/-^ cells. We bred *Cgas^-/-^, Sting^gt/gt^,* and *Tlr4*^-/-^ to *Casp1^-/-^Casp11*^-/-^ mice to generate triple mutants (*Casp1^-/-^Casp11^-/-^Cgas^-/-^, Casp1^-/-^Casp11^-/-^Sting^gt/gt^*, and *Casp1^-/-^Casp11^-/-^Tlr4^-/-^)* and then infected BMDMs from these mice. As expected, due to their inability to induce pyroptosis, all three triple mutants exhibited reduced host cell death, similar to levels seen for *Casp1^-/-^Casp11^-/-^*double mutants (**Figure 2A**). We next measured IFN-I secretion and found that infected *Casp1^-/-^Casp11^-/-^Tlr4^-/-^*cells maintained increased levels of IFN-I, similar to *Casp1^-/-^ Casp11^-/-^* cells (**Figure 2B**). In contrast, *Casp1^-/-^Casp11^-/-^Cgas^-/-^*and *Casp1^-/-^Casp11^-/-^Sting^Gt/Gt^*macrophages exhibited reduced IFN-I secretion, below levels of infected WT cells (**Figure 2B**). To determine whether the amount of IFN-I production correlated with an effect on bacterial growth, we also measured PFU over time. Mutation of *Cgas* or *Sting* in the *Casp1^-/-^Casp11^-/-^* background rescued growth of *R. parkeri* (**Figure 2C**) (for unknown reasons at 96 hpi, bacterial burdens were consistently lower in *Casp1^-/-^Casp11^-/-^Cgas^-/-^*and *Casp1^-/-^Casp11^-/-^Sting^gt/gt^* cells than in WT cells). In contrast, mutation of *Tlr4* in the *Casp1^-/-^Casp11^-/-^* background did not rescue bacterial growth. Together, these data in macrophages demonstrate that caspase-11 activation masks cGAS-dependent IFN-I production, and are consistent with the notion that pyroptosis prevents IFN-I production (Corrales et al., 2016; Liu et al., 2018). This observation suggests that, in cells where bacteria release DNA to activate cGAS, they also release LPS to activate caspase-11.

**Figure 2:**
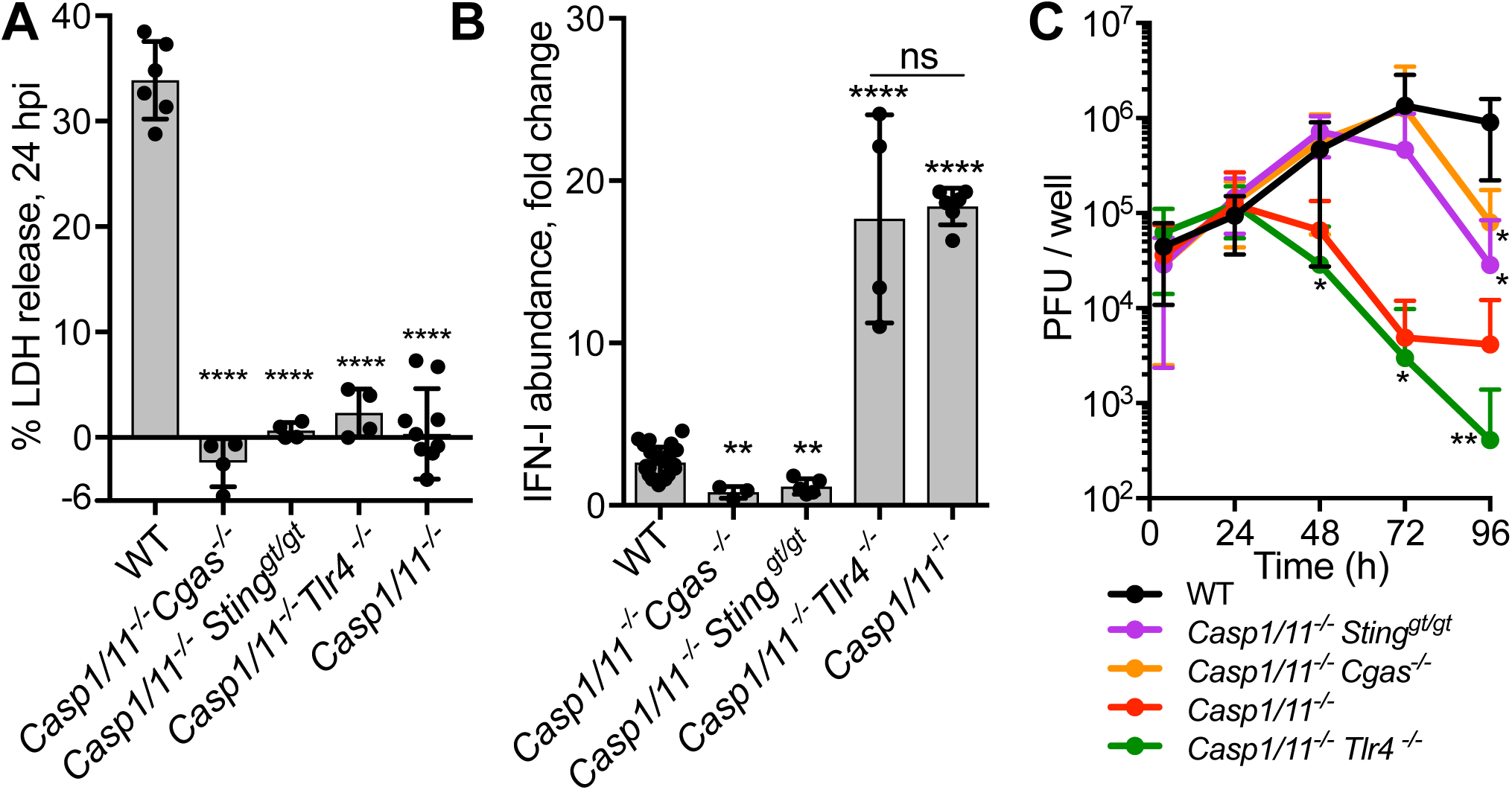
Pyroptosis masks cGAS-induced IFN-I. **A)** Quantification of host cell death during *R. parkeri* infection of BMDMs. LDH release was measured at 24 hpi upon *R. parkeri* infection of the indicated BMDMs at an MOI of 1. Statistical comparisons were made between each sample and WT B6 cells. Data are the compilation of at least three separate experiments and are expressed as means ± SEM. **B**) Measurement of IFN-I abundance in supernatants of infected BMDMs. The indicated BMDMs were infected with *R. parkeri* at an MOI of 1. Supernatants from infected cells were harvested at 24 hpi and used to stimulate a luciferase-expressing cell line and compared to uninfected cells. Statistical comparisons were made between each sample and infected WT cells. Data are a compilation of at least three separate experiments and are expressed as means ± SEM. **C**) Measurement of bacterial PFU in BMDMs over time. BMDMs were infected with *R. parkeri* at an MOI of 1 and PFU were measured over time. Statistical comparisons were made between each sample and WT B6 cells at each time point. Data are the compilation of at least three separate experiments and are expressed as means ± SEM. Statistical analyses for bacterial abundance in panel C were performed using a two-tailed Student’s T-test, where each sample at each time point was compared to the control. Statistical analyses for LDH assays and IFN-I abundance in panels A and B were performed using an ANOVA with multiple comparisons post-hoc test. *p<0.05, **p<0.01, ***p<0.001, ****p<0.0001, ns=not significant.

### IRF5-regulates antimicrobial genes and is required for IFN-I-dependent killing of *R. parkeri*

The host factors that restrict *R. parkeri* cytosolic growth downstream of IFN-I are poorly understood, in part because IFN-I upregulates hundreds of ISGs, complicating the ability to identify specific antimicrobial factors. To narrow the list of potential ISGs that restrict *R. parkeri* growth, we first tested whether a specific IRF was required for killing *R. parkeri* upon IFN-I treatment. We analyzed bacterial growth over time following bacterial infection of WT, *Irf1*^-/-^, *Irf3^-/-^Irf7*^-/-^, or *Irf5*^-/-^ mutant BMDMs and treatment with IFN-I. We observed that mutation of *Irf5* significantly rescued bacterial growth compared to WT cells, whereas mutation of *Irf1* or both *Irf3* and *Irf7* caused a modest increase in growth (**Figure 3A**). This suggested that genes specifically regulated by IRF5 restrict *R. parkeri*.

**Figure 3:**
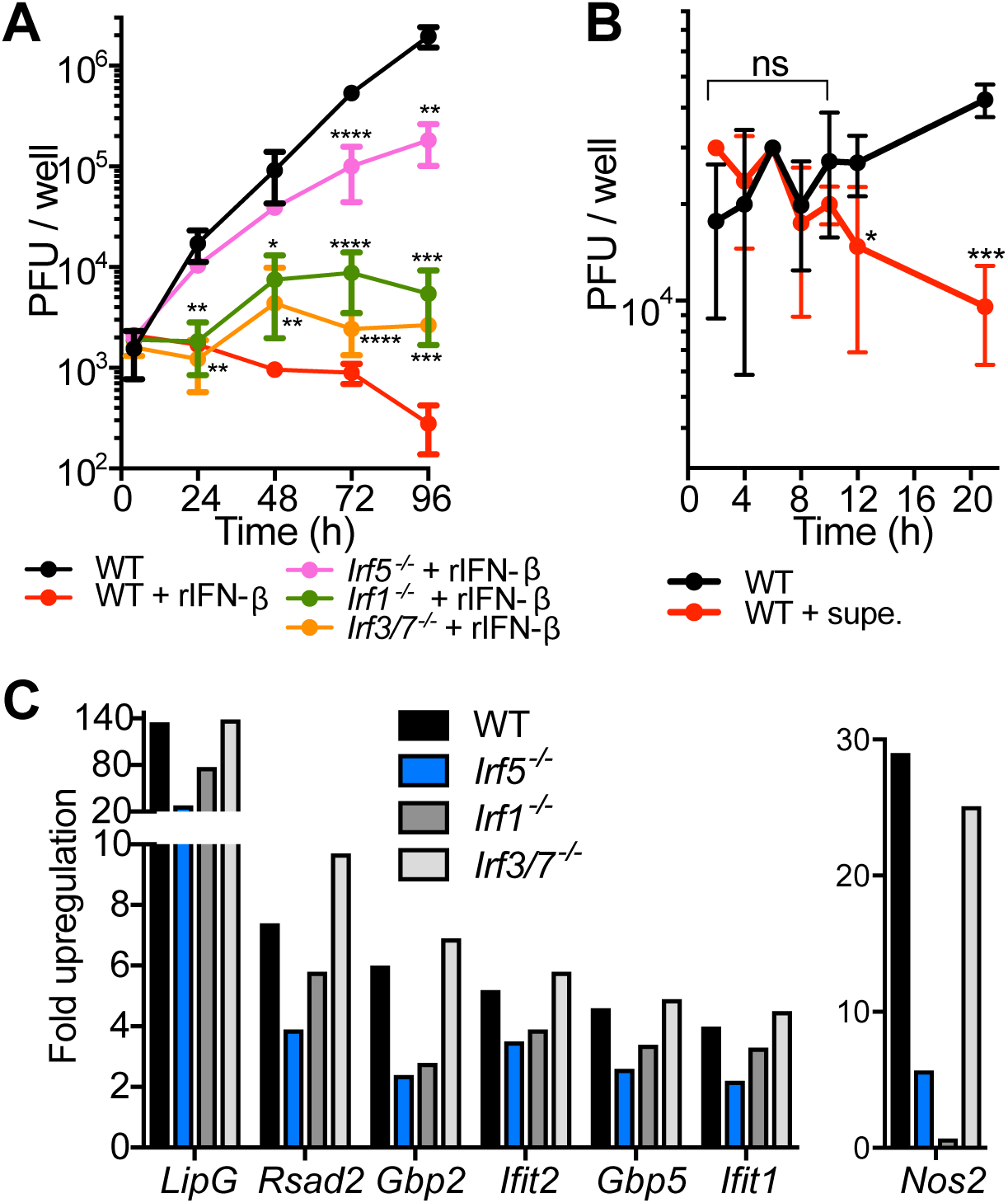
IRF5 is required for IFN-I-dependent restriction of *R. parkeri*. **A**) Measurement of *R. parkeri* PFU in BMDMs over time. Cells were infected at an MOI of 0.2, and PFU were measured over time. 10,000 units of recombinant IFN-β (rIFN-β) was added at 0 hpi. Statistical comparisons were made between each sample and untreated cells at each time point. Data are the compilation at least three separate experiments and are expressed as means ± SEM. **B**) Measurement of bacterial PFU in BMDMs. Cells were infected at an MOI of 1, and PFU were measured over time. Statistical comparisons were made between each sample and WT B6 cells at each time point. Data are the compilation at least three separate experiments and are expressed as means ± SEM. “Supe” indicates 200 µl of supernatant collected at 24 hpi from *Casp1^-/-^Casp11^-/-^* BMDMs infected with *R. parkeri* at an MOI of 1. All statistical analyses were performed using a two-tailed Student’s T-test. *p<0.05, **p<0.01, ***p<0.001, ****p<0.0001, ns=not significant. **C**) Transcript abundance of antimicrobial genes upregulated by IFN-I. BMDMs were infected at an MOI of 2.3, treated with 10,000 U recombinant IFN-β, and RNA was harvested at 12 hpi. High-throughput RNA-sequencing was then performed. The regulation of 7 candidate anti-rickettsial genes is shown. Genes are ordered from highest to lowest in terms of upregulation by IFN-β in WT cells, with the exception of *Nos2*, which is shown separately because it was more highly regulated by IRF1 than IRF5. Each data set was normalized to *R. parkeri-*infected WT, IFN-β-untreated BMDMs.

To identify genes significantly and specifically regulated by IRF5, we performed high-throughput RNA-sequencing (RNAseq) on the different WT, *Irf1*^-/-^, *Irf3^-/-^Irf7*^-/-^, or *Irf5*^-/-^ mutant BMDMs upon infection and IFN-I treatment. To limit the identification of genes not directly responsible for bacterial killing, we isolated RNA at 12 hpi, the earliest time when bacterial killing was observed (**Figure 3B**). Analysis of the RNAseq data revealed that 136 genes were upregulated >4.0-fold in infected IFN-I-treated cells when compared to infected untreated cells (**Table S1**). Among these, 82 were not as highly upregulated (>1.5-fold difference) in infected IFN-I-treated *Irf5^-/-^* cells when compared to infected IFN-I-treated WT cells, suggesting that many were regulated by IRF5. We then compared the expression of these genes between infected IFN-I-treated *Irf5*^-/-^ and *Irf1^-/-^* and *Irf3^-/-^Irf7*^-/-^ cells, and 36 had higher expression in *Irf1^-/-^* and *Irf3^-/-^Irf7*^-/-^ cells than in *Irf5^-/-^* cells. From the analysis of the RNA-seq data, we conclude that 36 genes are significantly and specifically upregulated by IRF5 during *R. parkeri* infection and IFN-I treatment (**Table S1**).

Many of the 36 genes encode known anti-viral and anti-bacterial proteins, including *Gbp2*, *Gbp5*, *Rsad2* (encoding Viperin), *Ifit1*, *Ifit2, Mx1,* and *Mx2* (**Figure 3C**). The RNA-seq also identified *Nos2*, encoding inducible nitric oxide synthase (iNOS), which is required for controlling *Rickettsia* spp. infection (Feng and Walker, 2000; Turco and Winkler, 1993; Turco et al., 1998; Walker et al., 1997), although its expression was also dependent on IRF1 (**Figure 3C**, right side). Together, these data demonstrate that IRF5 is critical for controlling *R. parkeri* upon IFN-I signaling, perhaps due to upregulating ISGs such as the GBPs and iNOS.

### GBPs and nitric oxide contribute to restricting *R. parkeri* growth in macrophages

We next assessed whether the IRF5-regulated genes that have previously known antimicrobial activity are important for inhibiting *R. parkeri* growth in the presence or absence of IFN-I. We derived BMDMs from the femurs of mice lacking *Ifit1, Ifit2, Rsad2, LipG* (the gene most highly upregulated by IFN-I), and the chromosome 3 cluster of *Gbps* (*Gbp^chr3^*) (Yamamoto et al., 2012), which includes *Gbp2*, *Gbp5* and three other *Gbp* genes. Bacterial growth was then measured upon IFN-I treatment. In IFN-I-treated *Gbp^chr3-/-^* macrophages, bacterial growth was increased 3-to-5-fold compared with that in WT IFN-I-treated macrophages (**Figure 4A**). No change in bacterial growth was observed in the other mutant macrophages upon IFN-I treatment (**Figure S2B**). When pyroptosis was assessed by measuring LDH-release, *Gbp^chr3^-*deficient macrophages showed dramatically reduced cell death upon infection, whereas the other mutant cells did not (**Figure 4B**). *Gbp^chr3-/-^* macrophages did not exhibit increased IFN-I production (**Figure 4C**), consistent with the notion that the GBPs are required for release of bacterial LPS as well as DNA. These data suggested that GBPs contribute to both IFN-I-dependent and IFN-I-independent killing of *R. parkeri*. Next, to assess the role for nitric oxide during *R. parkeri* infection, we treated WT- and *Gbp^chr3^*- infected cells with IFN-I and the specific iNOS inhibitor L-NIL and monitored bacterial growth. Upon iNOS inhibition, bacterial growth was increased in both WT and *Gbp^chr3-/-^*cells (**Figure 4D**, **4E**). We also measured bacterial growth in IFN-I-treated *Nos2^-/-^*BMDMs, but did not observe any rescue in mutant versus WT cells (**Figure S4A**), but this result is complicated by the fact that iNOS is a negative regulator of the NLRP3 inflammasome (Hernandez-Cuellar et al., 2012; Mishra et al., 2013) and we observed more host cell death in *Nos2^-/-^*BMDMs compared to WT cells (**Figure S4B**). All together, these experiments reveal that the GBPs and perhaps nitric oxide are IRF5-regulated genes that non-redundantly restrict *R. parkeri* growth in macrophages upon IFN-I treatment.

**Figure 4:**
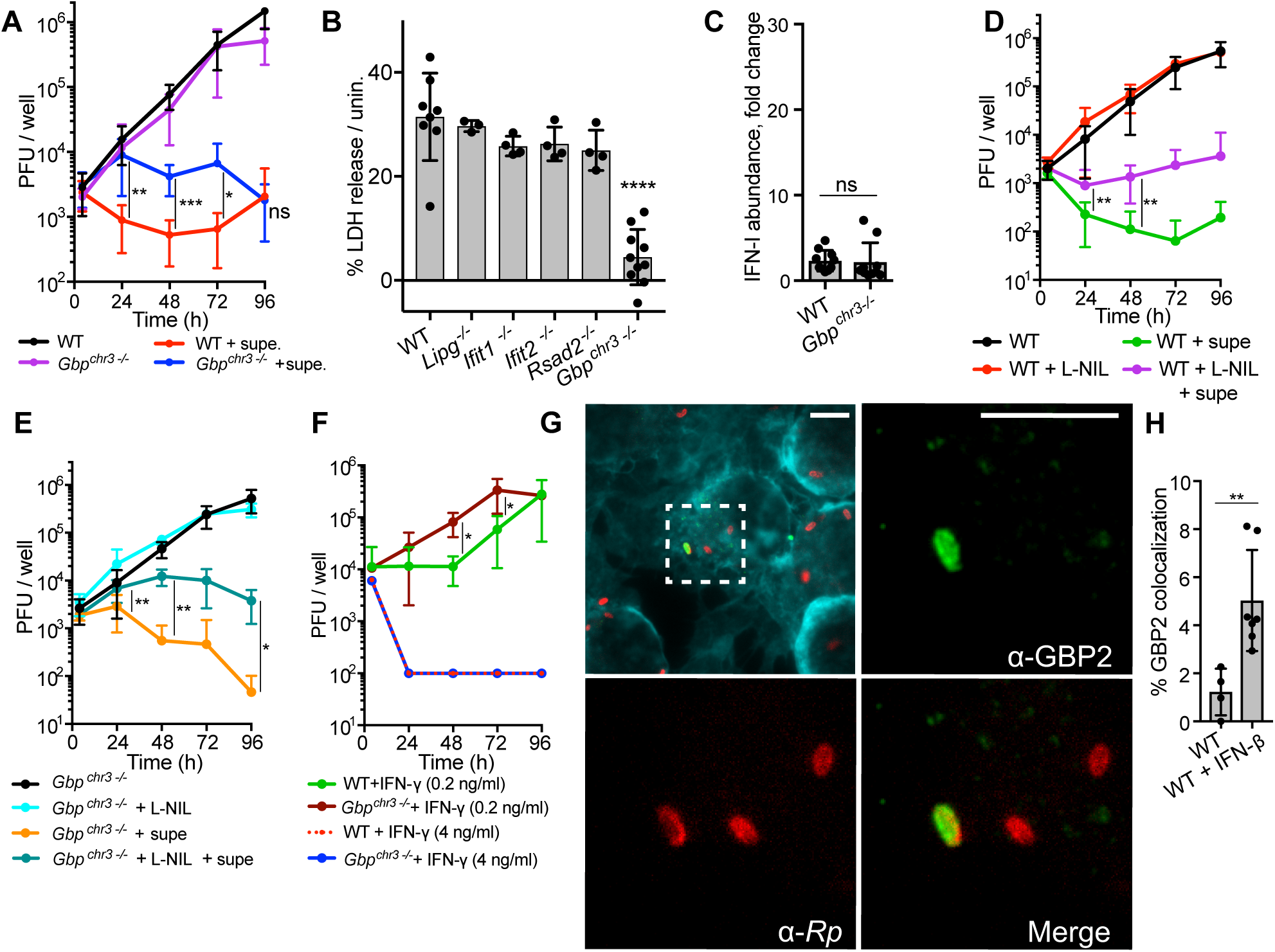
The GBPs contribute to IFN-I-dependent and -independent inhibition of *R. parkeri* growth. **A**) Measurement of *R. parkeri* PFU in BMDMs. BMDMs were infected with *R. parkeri* at an MOI of 0.2 and PFU were measured over time. “Supe” signifies 500 µl supernatant from infected *Casp1^-/-^Casp11^-/-^* cells, which was added at 0 hpi. Statistical comparisons were made between each sample and untreated cells at each time point. Data represent at least three separate experiments and are expressed as means ± SEM. **B**) Quantification of host cell death. LDH release was measured at 24 hpi upon *R. parkeri* infection at an MOI of 1. Statistical comparisons were made between each sample and WT B6 cells. Data are the compilation at least three separate experiments and are expressed as means ± SEM. **C)** Measurement of IFN-I abundance in supernatants of infected BMDMs. The indicated BMDMs were infected with *R. parkeri* at an MOI of 1. Supernatants from infected cells were harvested at 24 hpi and used to stimulate a luciferase-expressing cell line and compared to uninfected cells. Data are a compilation of at least five separate experiments. **D,E,F**) Measurement of bacterial PFU in BMDMs. Cells were infected at an MOI of 0.2, and PFU were measured over time. “Supe” signifies 500 µl supernatant from infected *Casp1^-/-^Casp11^-/-^* cells, which was added at 0 hpi. L-NIL was resuspended in water and added to a final concentration of 1 mM at T=0. IFN-γ was added at 0 hpi. Statistical comparisons were made between each sample and untreated cells at each time point. Data are the compilation of at least three separate experiments and are expressed as means ± SEM. **G**) A representative image of GBP2 localization to the surface of *R. parkeri.* WT BMDMs were infected at an MOI of 1, and were imaged at 3 hpi using 100x confocal immunofluorescence microscopy. Cyan staining is phalloidin; green staining is α-GBP2 (ProteinTech); red staining is α-*Rickettsia.* The red and green channels of the indicated white box are increased in size and shown on the right and bottom left. The scale bar is 5.6 μm. **H**) Quantification of GBP2 localization to the surface of *R. parkeri.* WT BMDMs were infected at an MOI of 1, and were imaged at 3 hpi using 100x confocal immunofluorescence microscopy. For IFN-β treated cells, BMDMs were treated overnight with 100 U recombinant IFN-β. Each data point represents an individual experiment, and each experiment consists of at least 10 separate images, and each image contained approximately 20 bacteria. Statistical analyses for panels A, C, D, E, F, and H were performed using a two-tailed Student’s T-test, where each sample at each time point was compared to the control. Statistical analyses for LDH assays in panel B were performed using an ANOVA with multiple comparisons post-hoc test. *p<0.05, **p<0.01, ***p<0.001, ****p<0.0001, ns=not significant.

GBP-mediated killing of other pathogens has also been examined downstream of IFN-γ treatment (Pilla-Moffett et al., 2016; Yamamoto et al., 2012). To assess the role of GBPs in IFN-γ- dependent restriction of *R. parkeri*, bacterial growth was measured in WT and *Gbp^chr3-/-^* cells in the presence of different concentrations of IFN-γ. At lower concentrations of IFN-γ (0.2 ng/ml), we observed a small but significant rescue in bacterial growth in *Gbp^chr3-/-^* versus WT cells (**Figure 4F**). However, higher concentrations of IFN-γ (4 ng/ml) ablated *R. parkeri* growth in both WT and *Gbp^chr3-/-^* BMDMs (**Figure 4F**). This revealed that IFN-γ potently stimulates macrophages to kill *R. parkeri* by a mechanism that partially depends on GBPs.

The GBPs have been observed to localize to the surface of intracellular pathogens at steady-state conditions and upon the addition of interferons (Liu et al., 2018; Mitchell and Isberg, 2017; Yamamoto et al., 2012). To determine if the GBPs localized to the surface of *R. parkeri,* we performed immunofluorescence microscopy using a GBP2-specific antibody. GBP2 localized to the surface of ∼1% of bacteria in untreated WT cells at 3 hpi, and ∼5% of total bacteria in IFN-I treated cells (**Figure 4G, H**). No colocalization was observed in infected *Gbp^chr3-/-^* macrophages, demonstrating that the antibody is specific for the GBPs (**Figure S3**). Together, these data demonstrate that the GBPs localize to *R. parkeri* and restrict bacterial growth at steady-state conditions, which is enhanced by IFN-I.

### In spleens, the inflammasome antagonizes the anti-rickettsial effects of IFN-I

Our data from macrophages suggested a model for *R. parkeri* intracellular survival whereby inflammasome activation antagonizes the anti-rickettsial activity of IFN-I. To determine whether this pathway is important *in vivo*, we analyzed the role for the inflammasome, IFN-I, and IFN-γ in spleens of infected mice, where *Rickettsia* spp. proliferate upon systemic infection (Feng et al., 1994). In agreement with the data from macrophages *in vitro*, IFN-β mRNA was significantly upregulated in spleens of infected *Casp1^-/-^Casp11^-/-^* mice when compared to WT mice, as measured by qPCR (**Figure 5A**). To determine if the increased IFN-β transcript abundance was dependent on cGAS, we also measured IFN-β mRNA in infected *Casp1^-/-^Casp11^-/-^Cgas^-/-^*mice, and observed a reduction in IFN-β transcript abundance (**Figure 5A**). This demonstrated that the inflammasome antagonizes cGAS-induced IFN-I *in vivo*, as it does *in vitro*.

**Figure 5:**
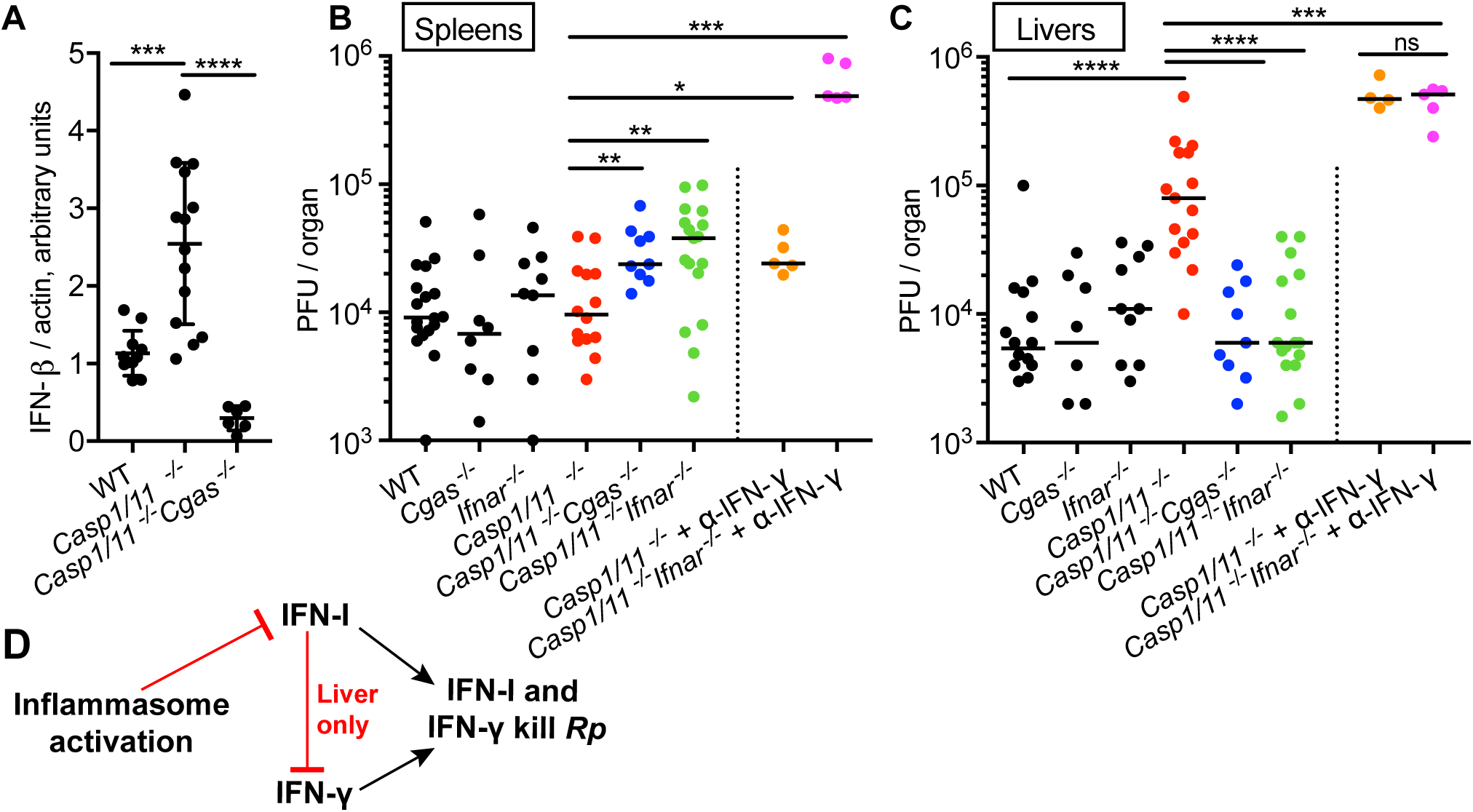
In mice, inflammasome activation antagonizes IFN-I production, leading to tissue-specific effects on *R. parkeri* burdens. **A**) Measurement of IFN-β transcripts in infected mice. C57BL/6 mice were infected i.v. with 10^7^ *R. parkeri,* and qPCR was used to analyze IFN-β and actin transcript abundance at 72 hpi in spleens. IFN-β was normalized to actin to determine relative copy number per mouse spleen. Data are the combination of at least 3 separate experiments. **B**) Measurement of bacterial burdens in mouse spleens. Mice were infected i.v. with 10^7^ *R. parkeri,* and bacterial burdens were determined in spleens at 72 hpi via plaque assay. Colors were arbitrarily assigned to distinguish between genotypes. Data are the combination of at least 3 separate experiments. Statistical analyses were performed using Mann-Whitney U test. Bars denote medians. **C**) Measurement of bacterial burdens in mouse livers. Mice were infected i.v. with 10^7^ *R. parkeri* and bacterial burdens were determined in livers at 72 hpi via plaque assay. For α-IFN-γ (BioLegend), mice were injected i.v. with 300 µl at 30 minutes postinfection (mpi), 200 µl at 24 hpi, and 200 µl at 48 hpi, totaling 800 µl (0.8 µg antibody). Colors are used to distinguish between genotypes. Data are the combination of at least 3 separate experiments, with the exception of experiments using the α-IFN-γ antibody, which are a combination of 2 separate experiments. Statistical analysis performed for qPCR experiments in panel A used a two-tailed Student’s T Test. Statistical analyses for *in vivo* experiments in panels B and C were performed using a Mann-Whitney U test. Bars denote medians. *p<0.05, **p<0.01, ***p<0.001, ****p<0.0001, ns=not significant. **D**) Textual summary of results from *in vivo* infections.

To determine if the increased IFN-I observed in *Casp1^-/-^Casp11^-/-^* mice restricted *R. parkeri,* we compared bacterial burdens between *Casp1^-/-^Casp11^-/-^*and *Casp1^-/-^Casp11^-/-^Ifnar^-/-^* mice. The triple *Casp1^-/-^Casp11^-/-^Ifnar^-/-^*mutant showed a significantly increased bacterial burden in comparison to the double *Casp1^-/-^Casp11^-/-^* mutant (**Figure 5B**). To ascertain if the increased bacterial growth was due to cGAS-dependent IFN-I, we also measured bacterial burdens in *Casp1^-/-^Casp11^-/-^Cgas^-/-^*mice, and similarly observed increased bacterial burdens compared with the *Casp1^-/-^Casp11^-/-^* mutant (**Figure 5B**). It remains unclear why spleens of *Casp1^-/-^Casp11^-/-^* mice, which produce more IFN-β, do not have reduced bacterial burdens when compared to WT (as in macrophages); nevertheless, these data reveal that the mechanisms of *R. parkeri* growth in spleens is similar to macrophages *in vitro*, where bacterial activation of the inflammasome limits the antimicrobial effects of cGAS-induced IFN-I.

In macrophages, we also observed a role for IFN-γ in restricting *R. parkeri* growth; however, the role for IFN-γ and its relationship to the inflammasome during *Rickettsia* infection *in vivo* is unknown. We therefore next assessed if IFN-γ restricted *R. parkeri* growth in the spleens of infected mice. Neutralization of IFN-γ using an anti-IFN-γ antibody increased bacterial burdens in *Casp1^-/-^Casp11^-/-^* mice 2.5-fold when compared to untreated *Casp1^-/-^Casp11^-/-^* mice, suggesting that IFN-γ restricts bacterial growth (**Figure 5B**, right side of dotted line). To reveal if IFN-γ and IFN-I both control infection in the spleen, we neutralized IFN-γ in infected *Casp1^-/-^Casp11^-/-^Ifnar^-/-^*mice. This increased bacterial burdens 51-fold over untreated *Casp1^-/-^Casp11^-/-^* mice (**Figure 5B**, right side of dotted line). These data demonstrate that in spleens of WT mice, inflammasome activation limits protective IFN-I production, and that IFN-I and IFN-γ both contribute to protection.

### In livers, the inflammasome antagonizes IFN-I, which in turn antagonizes the anti-bacterial effects of IFN-γ

We also measured bacterial burdens in the livers, where *Rickettsia* and other bacterial pathogens also reside during infection (Feng et al., 1994; Rayamajhi et al., 2010). In contrast to what was observed in spleens, infected *Casp1^-/-^Casp11^-/-^* mice had 15-fold higher bacterial burdens than WT mice in the liver (**Figure 5C**). Also in contrast to what was observed in spleens, the bacterial burden in infected *Casp1^-/-^Casp11^-/-^Ifnar^-/-^*mice, as well as *Casp1^-/-^Casp11^-/-^Cgas^-/-^*mice, was indistinguishable from WT (**Figure 5C**), suggesting that the 15-fold increase observed in *Casp1^-/-^Casp11^-/-^*mice was dependent on IFN-I. We hypothesized that this effect might be due to IFN-I antagonizing the anti-bacterial effects of IFN-γ, as it was previously demonstrated that IFN-I antagonizes the effects of IFN-γ in the liver during *L. monocytogenes* infection (Rayamajhi et al., 2010). To test whether IFN-I antagonized the anti-rickettsial effects of IFN-γ, infected *Casp1^-/-^Casp11^-/-^*and *Casp1^-/-^Casp11^-/-^Ifnar^-/-^* mice were treated with the IFN-γ neutralizing antibody and bacterial burdens were measured. The difference in bacterial burdens between these mice was erased upon IFN-γ neutralization (**Figure 5C**, right), demonstrating that IFN-I was responsible for antagonizing IFN-γ. Together, these results suggest that the inflammasome protects against *R. parkeri* infection in the liver, in part by limiting IFN-I antagonism of IFN-γ (**Figure 5D**).

### Both IFN-I and IFN-γ are required to control *R. parkeri* infection in mice

Because we observed that bacterial burdens in organs increased when both IFN-I and IFN-γ were removed, we next tested whether these cytokines were protective at the whole animal level. We infected C57BL/6 mice deficient for both IFN-I and IFN-γ receptors (*Ifnar^-/-^Ifngr^-/-^*) via the intravenous route. Strikingly, whereas WT, *Ifnar^-/-^,* and *Ifngr^-/-^* mice survived an infectious dose of 10^7^ bacteria with no severe symptoms, the majority of *Ifngr^-/-^Ifnar^-/-^* mice succumbed within 6 d (**Figure 6A**), and lost body weight and had reduced body temperature over time (**Figure 6B,C**). Symptoms of infection were dose-dependent. At a dose of 10^6^ bacteria, mice lost weight, their temperature decreased, and many succumbed to infection, although some eventually recovered to their original body weight and temperature. At dose of 10^5^ bacteria, none of the mice succumbed to infection, but the animals lost weight and their body temperature decreased, which then recovered to pre-infection levels. We also performed these experiments using AG129 mice, which are similarly mutated for *Ifngr* and *Ifnar*, but are in a different genetic background. These mice were also highly susceptible to infection with 10^7^ bacteria, with 100% fatality by 6 dpi (**Figure 6D**). Together, these data demonstrate that both IFN-I and IFN-γ potently control *R. parkeri* growth in animals. The discovery of the sensitivity of this mouse genotype reveals it as robust animal model for further investigations into pathogenesis of the SFG *Rickettsia*.

**Figure 6:**
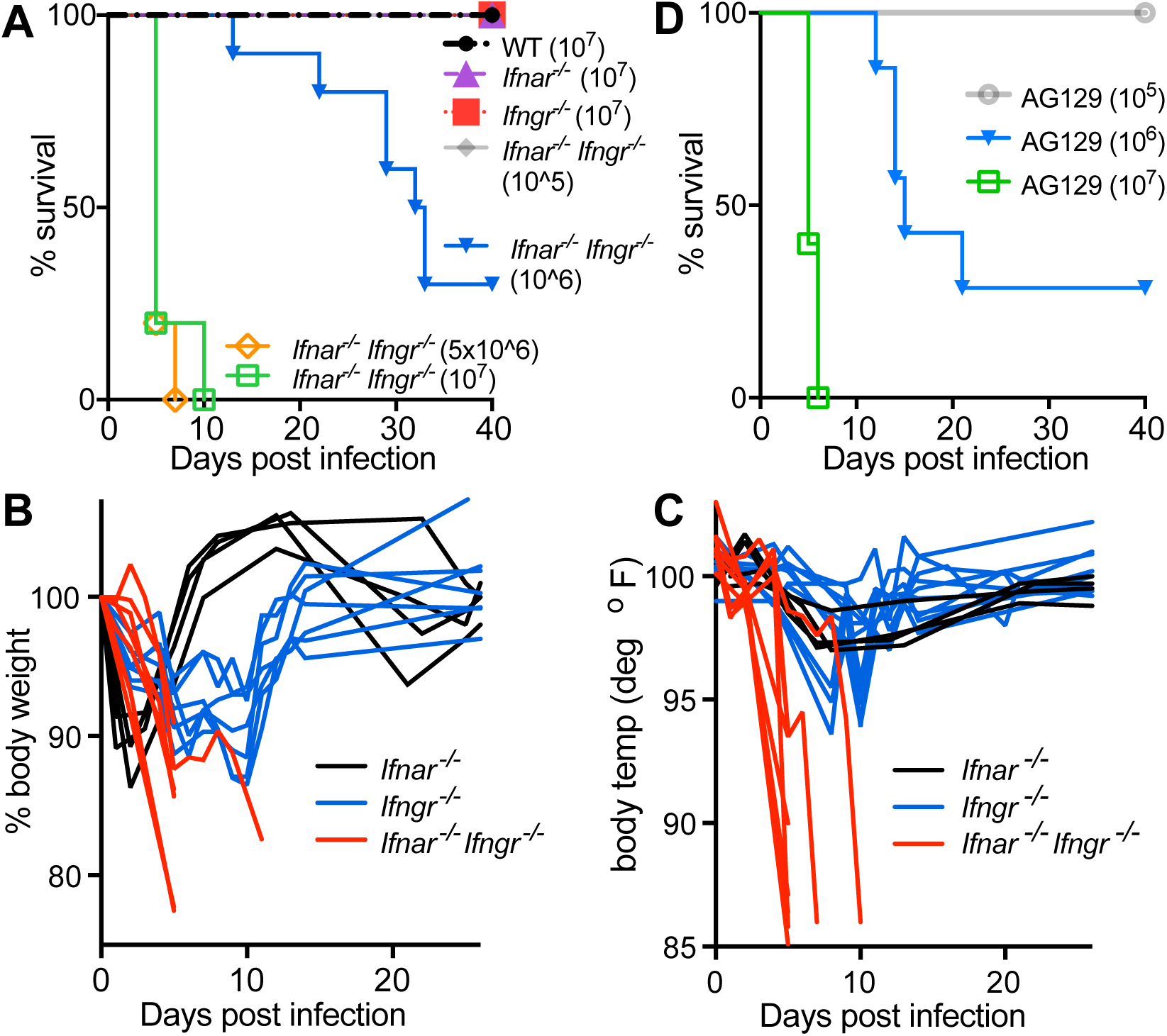
Mice mutated for both *Ifnar* and *Ifngr* succumb to *R. parkeri* infection. **A**) Survival of mice lacking interferon signaling. Mice in the C57BL/6 background were infected i.v. with the indicated concentrations of *R. parkeri,* and survival was measured over time. Mice were euthanized if their core body temperature fell below 90° F or if they exhibited symptoms of severe infection, such as severe hunching, scruffing, or inability to move around the cage. Infections from at least 6 mice are shown for each genotype. Data are the combination of at least three separate experiments. **B**) Measurement of mouse weight during infection. Mice were infected with 10^7^ *R. parkeri* and weight was measured every 24 hpi. Data are normalized to the weight starting at the initial time of infection. **C**) Measurement of mouse core body temperature during infection. Mice were infected with 10^7^ *R. parkeri* and body temperature was measured every 24 hpi, using a rectal thermometer. **D**) Survival of mice lacking interferon signaling. Mice of the 129 genotype lacking IFN-I and IFN-γ receptors (AG129) were infected i.v. with the indicated concentrations of *R. parkeri* and mouse health was measured over time. Mice were euthanized if their core body temperature fell below 90° F or if they exhibited symptoms of severe infection. At least 6 mice were used for each condition. Data are the combination of at least 3 separate experiments.

## Discussion

Avoiding innate immunity is critical for microbial pathogens to survive and cause disease. Whereas some facultative pathogens that inhabit the cytosol resist killing by IFN-I and avoid the inflammasome, the innate immune response to obligate intracellular bacteria has remained largely unexplored. We report the unexpected discovery that the innate immune response to the obligate intracellular human pathogen *R. parkeri* is distinctive among other cytosolic pathogens, as the bacteria are sensitive to IFN-I-mediated killing, but avoid stimulating a robust IFN-I response by exploiting the inherent trade-off between inflammasome activation and IFN-I production.

We observed that *R. parkeri* were restricted by IFN-I in macrophages and in mice. Furthermore, IFN-I much more potently restricted *R. parkeri* in macrophages than was previously seen for other *Rickettsia* spp. in immortalized endothelial cells and fibroblasts (Colonne et al., 2011; Turco and Winkler, 1990). In contrast, facultative cytosolic pathogens such as *L. monocytogenes* are resistant to the killing effects of IFN-I in primary myeloid cells (Bauler et al., 2011; Woodward et al., 2010) and in mice (Auerbuch et al., 2004; Rayamajhi et al., 2010). Given their sensitivity to IFN-I, we propose that *Rickettsia* must have evolved mechanisms to avoid stimulating IFN-I production.

Our results suggest that one critical mechanism for avoiding IFN-I production is that *R. parkeri*-induces pyroptosis, which dampens IFN-I production, protecting the remaining bacterial population that has successfully infected other cells (**Figure 7**, left). In support of this conclusion, we observed that a subpopulation of *R. parkeri* activated caspase-11 and GSDMD to cause pyroptosis. Because pyroptosis is a rapid post-translational signaling event (Kayagaki et al., 2015), activation of this cell death pathway prohibits IFN-I production, which is slower as it requires transcription, translation, and secretion. We found that macrophages that are deficient for inflammasome signaling restricted *R. parkeri* because bacterial ligands instead activated robust IFN-I production via cGAS, leading to bacterial killing (**Figure 7**, right). We also observed that the GBPs are required for pyroptosis and for IFN-I production in the absence of host cell death. From these observations, we conclude that bacteriolysis is likely responsible for the release of LPS and DNA that stimulate caspase-11 and cGAS. In contrast with what we observed for *R. parkeri*, several facultative bacterial pathogens have evolved different mechanisms to avoid the inflammasome (Jorgensen and Miao, 2015), including modification LPS by *F. novicida* to avoid caspase-11 (Hagar et al., 2013; Wallet et al., 2016), downregulation of flagellin by *L. monocytogenes* to avoid NAIP5 (Miao et al., 2010; Shen and Higgins, 2006), and infrequent lysis by *F. tularensis* and *L. monocytogenes* to avoid AIM2 (Sauer et al., 2010; Ulland et al., 2010). We propose that inflammasome activation is a trade-off that allows for growth of a pathogen that is sensitive to IFN-I.

**Figure 7:**
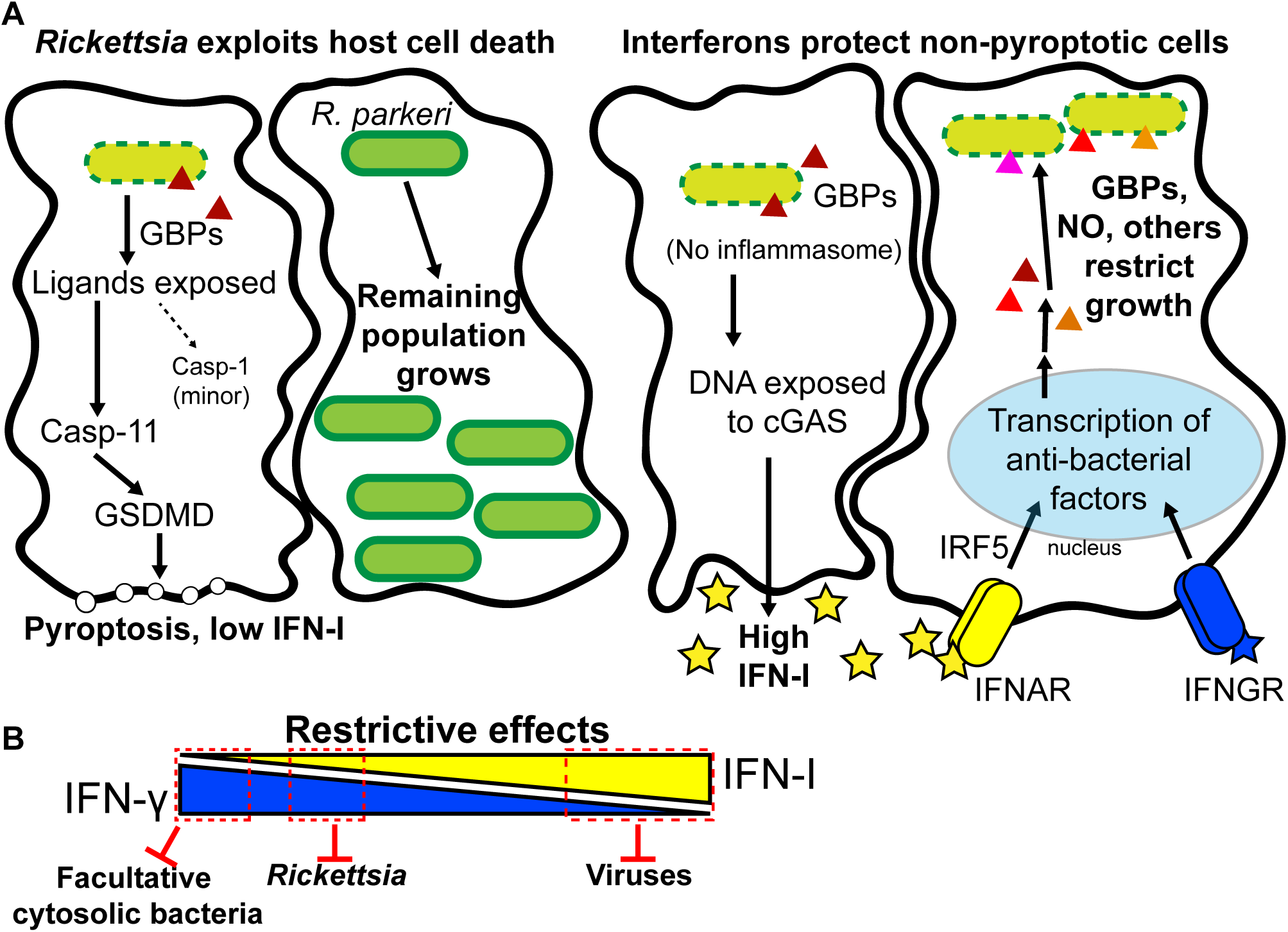
Inflammasome-mediated antagonism of IFN-I promotes intracellular growth of *R. parkeri*. **A**) Model depicting intracellular growth of *R. parkeri* in WT cells (left) or in cells lacking inflammasome signaling (right). **B**) Visual representation of the antimicrobial effects of interferons on cytosolic pathogens. Red zones indicate approximate restrictive effects of interferons.

Downstream of IFN-I production, we observed that the transcription factor IRF5 is critical for restriction of *R. parkeri* in macrophages *in vitro.* IRF5 in turn upregulates the expression of anti-microbial factors including the GBPs and iNOS, which protect against *R. parkeri*. These data agree with a previously established role for nitric oxide in protecting against infection by other *Rickettsia* spp. in mouse and human cells (Feng and Walker, 1993; 2000; Feng et al., 1994; Turco and Winkler, 1993; Woods et al., 2005), although it remains unclear if nitric oxide directly or indirectly kills these pathogens. These data also reveal an unappreciated role for the GPBs in rickettsial killing. We observed that iNOS and GBPs acted additively and non-redundantly, and only partially account for the killing effects of IFN-I. We suspect that additional IRF5-regulated genes contribute to controlling *R. parkeri* infection, and identifying these factors will be critical for understanding how interferons control infection by obligate pathogens.

Consistent with our observations in macrophages *in vitro*, we found that the inflammasome antagonizes IFN-I production *in vivo*. However, we observed tissue-specific differences in the spleen and liver with regard to the relative protective role for IFN-I. In the spleen, IFN-I protects against infection. These data are in agreement with previously known roles for the inflammasome in the spleen, where the major protective pathway elicited by the inflammasome is pyroptosis (Maltez et al., 2015). Our observations expand on these previous findings, by revealing that an inherent trade-off of pyroptosis is that it limits protective IFN-I in the spleen. However, in the liver, IFN-I enhances infection due to its effects of antagonizing IFN-γ. These observations are also in agreement with a previous study demonstrating that cytokine production is the more important facet of inflammasome activation in the liver (Maltez et al., 2015), and expand on these studies by demonstrating that the inflammasome also acts by antagonizing IFN-I. Our observations are also consistent with a study demonstrating that IFN-I antagonizes IFN-γ in the liver, as observed during *L. monocytogenes* infection (Rayamajhi et al., 2010). Moreover, our observations in both organs are consistent with earlier studies establishing a protective role for IFN-γ during *Rickettsia* infection *in vivo* (Feng et al., 1994; Li et al., 1987). Based on our findings that IFN-I and IFN-γ act differently in the spleen and liver, we speculate that interferons may pressure pathogens to evolve organ-specific tropisms to accommodate their intracellular lifestyle.

In keeping with our observations that IFN-I and IFN-γ restrict *R. parkeri* growth in macrophages *in vitro,* we observed that *Ifnar^-/-^Ifngr^-/-^* mice rapidly succumbed to infection. These findings demonstrate that both types of interferon play a critical role in protecting against *R. parkeri* infection. Our observations are also of strong practical importance, as this mouse exhibits a pathology that makes it a useful animal model for future studies examining the bacterial determinants that contribute to SFG *Rickettsia* pathogenesis and for evaluating the impacts of infection at the organ, tissue and cellular levels *in vivo*. Future studies will aim to characterize facets of infection including dissemination, vascular damage, as well as the innate immune response.

Our data suggest that obligate cytosolic pathogens such as *R. parkeri* may occupy an intermediate position in the spectrum between bacteria and viruses, both with regard to their dependency on the host and their interplay with the innate immune system (**Figure 7B**). For example, during infection with *R. parkeri* as well as with herpes simplex virus 1 or Vaccinia virus, activation of the inflammasome antagonizes the protective IFN-I response (Wang et al., 2017). Additionally, IRF5 is required for controlling *R. parkeri* and viral infection (Carlin et al., 2017; Proenca-Modena et al., 2016; Thackray et al., 2014). Lastly, as with *R. parkeri*, many viruses have increased lethality in *Ifnar^-/-^Ifngr^-/-^* mice (Milligan et al., 2017; Rossi et al., 2016), suggesting a non-redundant protective role for IFN-I and IFN-γ in both types of infection. These similarities are not true for facultative cytosolic bacterial pathogens such as *L. monocytogenes,* for which IFN-γ is significantly more protective than IFN-I (Rayamajhi et al., 2010). Our observations regarding the interaction between *R. parkeri* and innate immune pathways underscores the notion that obligate pathogens have evolved different strategies than facultative pathogens to fit their ecological niche. Their ability to tolerate inflammasome activation to avoid IFN-I is an evolutionary strategy that may be shared among obligate microbes, whose complete reliance on host cell metabolic and cellular pathways for survival may also dispose them to IFN-I sensitivity. Continued investigation of the interaction between obligate intracellular bacteria and innate immunity will enhance our understanding of how pathogens exploit host cell immunity to survive and to cause disease.

## Acknowledgements

We thank Dr. Mike Diamond (Washington University, St. Louis) for femurs from *Irf5^-/-^*, *Ifit1^-/-^*, and *Ifit2^-/-^* mice. We thank Dr. Daniel Rader (Pittsburg University) for femurs from *LipG^-/-^* mice. We thank Dr. Jörn Coers (Duke University) and Dr. Masahiro Yamamoto (Osaka Yamamoto) for femurs from *Gbp^chr3-/-^* mice. We thank Dr. Eva Harris (UC Berkeley) for AG129 mice (originally obtained from M. Aguet, Swiss Institute for Experimental Research, Epalinges, Switzerland). We thank Dr. Greg Barton (UC Berkeley) for helpful advice and for *Irf3^-/-^Irf7*^-/-^ mice and *Tnfrsf1a^-/-^Tnfrsf1b^-/-^* mice. We thank Neil Fischer for critical reading of this manuscript. P.E. was supported by postdoctoral fellowships from the Foundation Olle Engkvist Byggmästare, the Swedish Society of Medical Research (SSMF), and the Sweden-America Foundation. M.D.W. was supported by NIH/NIAID grants R01 AI109044, R21 AI109270, and R21 AI138550. R.E.V. is an HHMI investigator and is supported by NIH/NIAID grants AI075039 and AI063302.

## Author contributions

Conceptualization: T.P.B; Methodology: T.P.B., P.E.; Investigation: T.P.B., P.E., and R.C.; Visualization: T.P.B., P.E., R.E.V., and M.D.W.; Writing: Original Draft, T.P.B.; Writing – Review & Editing, T.P.B., P.E., R.E.V., and M.D.W.; Funding Acquisition, R.E.V. and M.D.W.; Resources, R.E.V. and M.W.D.; Supervision, R.E.V. and M.D.W.

## STAR Methods

### Contact for reagent and resource sharing

Further information and requests for reagents may be directed to and will be fulfilled by the Lead Contact Dr. Matthew D. Welch.

## EXPERIMENTAL MODEL AND SUBJECT DETAILS

### Bacterial strains

All *R. parkeri* used in this study were in the Portsmouth strain originally obtained from Christopher Paddock (Centers for Disease Control and Prevention) and were authenticated by whole genome sequencing (NCBI Trace and Short-Read Archive; Sequence Read Archive (SRA), accession number SRX4401164). To prepare the bacteria for infections, confluent T175 flasks of female African green monkey kidney epithelial Vero cells (authenticated by STR analysis) grown in DMEM (Gibco 11965-092) with glucose (4.5 g/L) supplemented with 2% fetal bovine serum (FBS, GemCell) were infected with 5 x 10^6^ *R. parkeri* per flask and rocked for 10 min at 37°C. Infected cells were scraped and collected at 5-6 dpi when ∼90% of cells were highly infected, as indicated by cell rounding. Scraped cells were centrifuged at 12,000 x G for 20 min at 4°C. Pelleted cells were then resuspended in K-36 buffer (0.05 M KH_2_PO_4_, 0.05 M K_2_HPO_4_, 100 mM KCl, 15 mM NaCl, pH 7) and transferred to a cold glass Dounce homogenizer. Bacteria were released from infected cells by repeated douncing (60 strokes). The dounced solution was then centrifuged at 200 x G for 5 min at 4°C to pellet host cell debris. The supernatant containing *R. parkeri* was overlaid on a 30% MD-76R (Merry X-Ray) solution. Gradients were then centrifuged at 18,000 rpm in an SW-28 ultracentrifuge swinging bucket rotor (Beckman/Coulter) for 20 min at 4°C to separate remaining host cells debris from the bacteria. Bacterial pellets were resuspended in brain heart infusion (BHI) media (BD, 237500) and stored at −80°C.

Titers were determined for *R. parkeri* stocks via plaque assays by serially diluting the bacteria in 6-well plates containing confluent Vero cells, which were plated ∼24 h prior. Plates were then spun for 5 min at 300 x G in an Eppendorf 5810R centrifuge. At 24 hpi, the media from each well was aspirated and the wells were overlaid with 4 ml/well DMEM with a final concentration of 5% FBS and 0.7% ultrapure agarose (Invitrogen, 16500-500). When plaques were visible at 5-6 dpi, an overlay of 0.7% agarose in DMEM containing 2.5% neutral red solution (Sigma, N6264) was added and plates incubated overnight until plaques were clearly visible. Plaques were then counted to determine bacterial concentrations.

*L. monocytogenes* strain 10403S was originally obtained from Dr. Daniel Portnoy (UC Berkeley).

### Animal experiments

Animal research using mice was conducted under a protocol approved by the UC Berkeley Institutional Animal Care and Use Committee (IACUC) in compliance with the Animal Welfare Act and other federal statutes relating to animals and experiments using animals. The UC Berkeley IACUC is fully accredited by the Association for the Assessment and Accreditation of Laboratory Animal Care International and adheres to the principles of the Guide for the Care and use of Laboratory Animals. Infectious disease studies were performed in a biosafety level 2 facility. All animals were maintained at the UC Berkeley campus and all infections were performed in accordance with the approved Welch lab Animal Use Protocol. Mice were age matched between 8 and 18 weeks old. Mice were selected for experiments based on their availability, regardless of sex. PFU data for each gender is reported in **Figure S5** at 48 and 72 hpi. All mice were healthy at the time of infection and were housed in microisolator cages and provided chow and water. Experimental groups were littermates of the same sex that were randomly assigned to experimental groups. All mice were phenotypically healthy at the time of the experiment, as determined by their weight, temperature, and movement around the cage. For experiments with mice mutated for *Ifnar* and *Ifngr*, as described in Figure 6, mice were immediately euthanized if they exhibited severe degree of infection, as defined by a core body temperature dropping below 90° F or lethargy that prevented the mouse from moving around the cage normally.

### Mouse genotyping

*Casp1^-/-^* (Rauch et al., 2017)*, Sting^gt/gt^* (Sauer et al., 2011)*, Cgas^-/-^* (Marcus et al., 2018)*, Gsdmd^-/-^* (Rauch et al., 2017)*, Irf5^-/-^* (Purtha et al., 2012), *Ifit1^-/-^* (Szretter et al., 2012), and *Ifit2^-/-^* (Fensterl et al., 2012), *lipG^-/-^* (Ishida et al., 2003), *Gbp^chr3^*^-/-^ (Yamamoto et al., 2012) mice were previously described. *Casp11^-/-^* (Wang et al., 1998)*, Irf1^-/-^* (Matsuyama et al., 1993), *Ifnar^-/-^* (Müller et al., 1994)*, Ifngr^-/-^* (Huang et al., 1993), *Ifnar^-/-^Ifngr^-/-^,* and WT mice were previously described and originally obtained from Jackson Laboratories. For genotyping, ear clips were boiled for 15 min in 60 µl of 25 mM NaOH, quenched with 10 µl tris-HCl pH 5.5, and 2 µl of lysate was used for PCR using SapphireAMP (Takara, RR350) and primers specific for each gene. Mutant mice were genotyped using the following primers: *Ifnar* forward (F): CAACATACTACAACGACCAAGTGTG; *Ifnar* WT reverse (R): AACAAACCCCCAAACCCCAG; *Ifnar* mutant R: ATCTGGACGAAGAGCATCAGG; WT *Casp1/11* F: CATGCCTGAATAATGATCACC; WT *Casp1/11* R: GAAGAGATGTTACAGAAGCC; *Casp1/11* mutant F: GCGCCTCCCCTACCCGG; *Casp1/11* mutant R: CTGTGGTGACTAACCGATAA; *Cgas* F: ACTGGGAATCCAGCTTTTCACT; *Cgas* R: TGGGGTCAGAGGAAATCAGC; WT *tlr4* F: CACCTGATACTTAATGCTGGCTGTAAAAAG; WT *tlr4* R: GGTTTAGGCCCCAGAGTTTTGTTCTTCTCA; *tlr4* mutant F: TGTTGCCCTTCAGTCACAGAGACTCTG; *tlr4* mutant R: TGTTGGGTCGTTTGTTCGGATCCGTCG; *Sting* F: GATCCGAATGTTCAATCAGC; *Sting* R: CGATTCTTGATGCCAGCAC; *Gsdmd* F: ATAGAACCCGTGGAGTCCCA; and *Gsdmd* R: GGCTTCCCTCATTCAGTGCT.

### Deriving bone marrow macrophages

To obtain bone marrow, male or female mice were euthanized and femurs, tibias, and fibulas were excised. Visible muscle and connective tissue were removed from the bones and the bones were sterilized with 70% ethanol. Bones were washed with BMDM media (20% HyClone FBS, 1% sodium pyruvate, 0.1% β-mercaptoethanol, 10% conditioned supernatant from 3T3 fibroblasts, in Gibco DMEM containing glucose and 100 U/ml penicillin and 100 ug/ml streptomycin) and ground using a mortar and pestle. Bone homogenate was passed through a 70 μm nylon Corning Falcon cell strainer (Thermo Fisher Scientific, 08-771-2) to remove particulates. Filtrates were centrifuged in an Eppendorf 5810R at 1,200 RPM (290 x G) for 8 min, supernatant was aspirated, and the remaining pellet was resuspended in BMDM media. Cells were then plated in non-TC-treated 15 cm petri dishes (at a ratio of 10 dishes per 2 femurs) in a volume of 30 ml BMDM media and incubated at 37° C. An additional 30 ml BMDM media was added 3 d later. At 7 d the media was aspirated and cells were incubated at 4°C with 15 ml cold PBS (Gibco, 1x pH 7.4, no ions) for 10 min. BMDMs were then scraped from the plate, collected in a 50 ml conical tube, and centrifuged at 1,200 RPM (290 x G) for 5 min. The PBS was then aspirated, cells were resuspended in BMDM media, and cell numbers were counted using trypan blue and a hemocytometer. For freezing, the pellet was resuspended in BMDM media with 30% FBS and 10% DMSO at 10^7^ cells/ml. 1 ml aliquots were frozen in a Styrofoam box at −80°C for one day and then moved to liquid nitrogen.

## METHOD DETAILS

### Mouse infections

For mouse infections, *R. parkeri* were prepared by diluting 30%-prep bacteria to 1 ml cold sterile PBS, centrifuging the bacteria at 12,000 x G for 1 min (Eppendorf 5430 centrifuge), and resuspended in cold sterile PBS to the desired concentration (either 5×10^7^ PFU/ml, 2.5×10^7^ PFU/ml, 5×10^6^ PFU/ml, or 5×10^5^ PFU/ml). Bacterial suspensions were kept on ice for the duration of the injections. For i.v. injections, mice were exposed to a heat lamp while in their cages for approximately 5 minutes and then each mouse was moved to a mouse restrainer (Braintree, TB-150 STD). The tail was sterilized with 70% ethanol and then 200 µl of bacterial suspensions were injected using 30.5 gauge needles into the lateral tail vein. For monitoring body temperatures of infected mice, mice were placed in a mouse restrainer, and a rodent rectal thermometer (BrainTree Scientific, RET-3) was used to measure temperature. For delivering the anti-IFN-γ antibody (BioLegend, 505847), mice were injected i.v. with 300 µl at 30 min p.i., 200 µl at 24 hpi, and 200 µl at 48 hpi, totaling 800 µl (0.8 µg antibody). Mice were monitored daily for clinical signs of disease, such as hunched posture, lethargy, or scruffed fur. Only mice lacking both interferon receptors exhibited such symptoms of disease, and if this occurred, mice were monitored daily for these symptoms, as well as for loss in body weight and temperature. If a mouse displayed severe symptoms of infection, as defined by a reduction in body temperature below 90°F or an inability to move around the cage normally, the animal was immediately and humanely euthanized using CO_2_ followed by cervical dislocation, according to IACUC-approved procedures.

For harvesting organs, mice were euthanized at the indicated pre-determined times and doused with ethanol. Mouse organs were extracted in a biosafety cabinet and deposited into 50 ml conical tubes (Falcon) containing 4 ml sterile cold PBS (Gibco 10010-023, no ions) for the spleen, and containing 8 ml cold sterile PBS for the liver. Organs were kept on ice and were each homogenized for approximately 10 s using an immersion homogenizer (Fisher, Polytron PT 2500E) at 22,000 RPM. Organ homogenates were spun at 290 x G for 5 min to pellet cell debris (Eppendorf 5810R centrifuge). 20 µl of organ homogenates were plated into confluent Vero cells (plated 48 h prior) in 12-well plates, and then serial diluted. The plates were then spun at 260 x G for 5 min at room temperature (Eppendorf 5810R centrifuge) and incubated at 33°C overnight. To reduce the possibility of contamination, organ homogenates were plated in duplicate and the second replicate was treated with 50 µg/ml carbenicillin (Sigma) and 200 µg/ml amphotericin B (Gibco). The next day, at approximately 16 hpi, the cells were gently washed by replacing the existing media with 1 ml DMEM containing 2% FBS. The media was then aspirated and replaced with 2 ml/well of DMEM containing 0.7% agarose, 5% FBS, and 200 µg/ml amphotericin B. When plaques were visible, at approximately 6 dpi, 1 ml of DMEM containing 0.7% agarose, 1% FBS, and 2.5% neutral red (Sigma) was added to each well. Plaques were counted 24 h later.

### Infections *in vitro*

Aliquots of BMDMs were thawed on ice, diluted into 9 ml of DMEM, centrifuged in an Eppendorf 5810R at 1,200 RPM (290 x G) for 5 minutes, and the pellet was resuspended in 10 ml BMDM media without antibiotics. The number of cells was counted using Trypan blue (Sigma, T8154) and a hemocytometer (Bright-Line), and 5 x 10^5^ cells were plated into 24-well plates. Approximately 16 h later, 30% prep *R. parkeri* were thawed on ice and diluted into fresh BMDM media to the desired concentration (either 10^6^ PFU/ml or 2×10^5^ PFU/ml). Media was then aspirated from the BMDMs, replaced with 500 µl media containing *R. parkeri,* and plates were spun at 300 G for 5 min in an Eppendorf 5810R. Infected cells were then incubated in a humidified CEDCO 1600 incubator set to 33°C and 5% CO_2_ for the duration of the experiment.

For measuring PFU, supernatants were aspirated from individual wells, and each well was gently washed twice with 500 µl sterile milli-Q-grade water. 1 ml of sterile milli-Q water was then added to each well and repeatedly pipetted up and down to lyse the host cells. Serial dilutions of lysates were added to confluent Vero cells in 12 well plates that were plated 24 or 48 h prior. Plates were then spun at 300 x G using an Eppendorf 5810R centrifuge for 5 min at room temperature and incubated at 33°C overnight. At ∼16 hpi, media was aspirated and replaced with 2 ml/well of DMEM containing 0.7% agarose and 5% FBS. When plaques were visible, at approximately 6 dpi, 1 ml of DMEM containing 0.7% agarose, 1% FBS, 200 µg/ml amphotericin B (Invitrogen, 15290-018), and 2.5% neutral red (Sigma) was added to each well. Plaques were counted 24 h later. For neutralizing IFN-I signaling, an ultra-LEAF-purified α-IFNAR-1 antibody (BioLegend, 127323) was added to a final concentration of 1 µg/ml at 0 hpi. For experiments with recombinant TNF-α, 200 ng was added to each well in a 24-well plate, and products from two different vendors were tested (BioLegend 575202 and Thermo PMC3014).

For collecting supernatant from *Casp1^-/-^Casp11^-/-^* cells, 5×10^5^ BMDMs in 24-well plates were infected with *R. parkeri* at an MOI of 1 and at 24 hpi supernatants were pooled and stored at −80°C. For adding the supernatant to infected BMDMs, either 200 or 500 µl of supernatant was removed at 20 mpi from the untreated cells and replaced with the supernatant from *Casp1^-/-^Casp11^-/-^* cells.

For infections with *L. monocytogenes,* cultures of *L. monocytogenes* strain 10403S were grown in 2 ml sterile-filtered BHI shaking at 37° until stationary phase (∼16 h). Cultures were centrifuged at 20,000 x G (Eppendorf 5430), the pellet was resuspended in sterile PBS (Gibco 10010-023), and diluted 100-fold in PBS. 10 µl of the diluted bacteria were then added to each well of a 24-well plate of BMDMs that were plated ∼16 h prior to infections at 5×10^5^ cells/well. Bacteria were also plated out onto Luria Broth agarose plates to determine the titer, which was determined to be ∼5 x 10^5^ bacteria / 10 µl, for an MOI of 1 (based on the ratio of bacteria in culture to number of BMDMs). Infected cells were incubated in a humidified 37° incubator with 5% CO_2_. 25 µg of gentamicin (Gibco 15710-064) was added to each well (final concentration 50 µg/ml) at 1 hpi. At 30 mpi, 2, 5, and 8 hpi, the supernatant was aspirated from infected cells, and cells were washed twice with sterile milli-Q water. Infected BMDMs were then lysed with 1 ml sterile water by repeated pipetting and scraping of the well. Lysates were then serially diluted and plated on LB agar plates, incubated at 37° overnight, and CFU were counted at ∼20 h later.

### High-throughput DNA sequencing

For high-throughput DNA sequencing, 5 x 10^5^ BMDMs were plated in 24-well plates and infected 16 h later with *R. parkeri* and treated with 10,000 activity units of recombinant IFN-β (PBL Cat # 12405-1). To determine the percentage of cells that were successfully infected, we analyzed the infected cells using immunofluorescence microscopy and observed that the multiplicity of infection (the average number of bacteria per host cell) was 2.3, and that 71% of cells were infected. At 12 hpi, cells were lysed and RNA was purified using an RNeasy purification kit (Qiagen). RNA quality was assessed using an Agilent 2100 Bioanalyzer, and all samples had RIN values above 8.0. Transcripts were selected using polyA selection (using Dynabeads mRNA Purification Kit, Invitrogen) and enzymatically fragmented as part of the Apollo library prep kits (Wafergen PrepX RNA library prep for Illumina). Libraries were constructed by using Apollo 324 (IntegenX Inc.), PCR-amplified, and multiplexed at the Functional Genomics Lab at the University of California, Berkeley (http://qb3.berkeley.edu/qb3/fgl/). The resulting libraries were sequenced at the Vincent J. Coates Genomics Sequencing Facility at the University of California, Berkeley using single-end reads, 50 base length, with the Hiseq 2000 Illumina platform. Sequence data were aligned to the *Mus musculus* C57BL/6 reference genome (reference assembly GCA_000001635.8) using CLC Genomics Workbench (Qiagen). Fold regulation of genes for each genotype was determined by referencing the sample uninfected, untreated BMDMs. Comparisons were then made between the sequencing results from the infected, IFN-I-treated WT and the *Irf* mutant genes. Each data set was composed of at least 55 million reads and 98.3% of the reads aligned with the reference. Genes with low abundance of reads (Reads Per Kilobase of transcript, per Million mapped reads, RPKM, of <10) in the infected WT BMDMs treated with IFN-I were excluded from the analysis. Sequencing data were uploaded to GEO, accession number GSE128211.

### Microscopy

For brightfield microscopy, images were captured using an IX71 Olympus microscope with a UCPLFLN 20x 0.7 NA objective, OptiMOS sCMOS camera (QImaging), and Micro-Manager software (Edelstein et al., 2014). For immunofluorescence microscopy, 2.5-5 x 10^5^ BMDMs were plated overnight in BMDM media in 24-well plates with sterile 12 mm coverslips (Thermo Fisher Scientific, 12-545-80). Infections were performed as described above. At the indicated times post-infection, coverslips were washed once with PBS and fixed in 4% paraformaldehyde (Ted Pella Inc., 18505, diluted in 1 x PBS) for 10 min at room temperature. Coverslips were then washed 3 times in PBS. Coverslips were washed once in blocking buffer (1 x PBS with 2% BSA) and permeabilized with 0.5% triton X-100 for 10 min. Coverslips were incubated with primary and secondary antibodies diluted in 2% BSA in PBS, each for 30 min at room temperature. *R. parkeri* were detected using mouse anti-*Rickettsia* 14-13 (originally from Dr. Ted Hackstadt, NIH/NIAID Rocky Mountain Laboratories). GBP2 was detected with anti-GBP2 (ProteinTech, 11854-1-AP; Research Resource Identifier AB_2109336). Nuclei were stained with DAPI, and actin was stained with Alexa-568 phalloidin (Life Technologies, A12380). Secondary antibodies were Alexa-405 goat anti-mouse (A31553) and Alexa-488 goat anti-rabbit (A11008). Coverslips were mounted in Prolong mounting media (Invitrogen). Samples were imaged with the Nikon Ti Eclipse microscope with a Yokogawa CSU-XI spinning disc confocal with 60X and 100X (1.4 NA) Plan Apo objectives, and a Clara Interline CCD Camera (Andor Technology) using MetaMorph software (Molecular Devices). Rendered Z-stacks were used for quantifications. Images were processed using FIJI (Schindelin et al., 2012) and brightness and contrast adjustments were applied to entire images. Images were assembled using Adobe Illustrator. Representative images are a single optical section, in which most or all bacteria were in the focal plane.

### *In vitro* assays

For LDH assays, 60 µl of supernatant from wells containing BMDMs was collected into 96-well plates. 60 µl of LDH buffer was then added. LDH buffer contained: 3 µl of “INT” solution containing 2 mg/ml tetrazolium salt (Sigma I8377) in PBS; 3 µl of “DIA” solution containing 13.5 units/ml diaphorase (Sigma, D5540), 3 mg/ml β-nicotinaminde adenine dinucleotide hydrate (Sigma, N3014), 0.03% BSA, and 1.2% sucrose; 34 µl PBS with 0.5% BSA; and 20 µl solution containing 36 mg/ml lithium lactate in 10 mM Tris HCl pH 8.5 (Sigma L2250). Supernatant from uninfected cells and from cells completely lysed with 1% triton X-100 (final concentration) were used as controls. Reactions were incubated at room temperature for 20 min prior to reading at 490 nm absorbance using an Infinite F200 Pro plate reader (Tecan). Values for uninfected cells were subtracted from the experimental values, divided by the difference of triton-lysed and uninfected cells, and multiplied by 100 to obtain percent lysis. Each experiment was performed and averaged between technical duplicates and biological duplicates.

For the IFN-I bioassay experiments, 5 x 10^4^ 3T3 cells containing an interferon-sensitive response element (ISRE) fused to luciferase (Jiang et al., 2005; McWhirter et al., 2009) were plated per well into 96-well white-bottom plates (Greneir 655083) in DMEM containing 10% FBS, 100 U/ml penicillin and 100 µg/ml streptomycin. 24 h later, media was replaced and confluent cells were treated with 2 µl of supernatant harvested from BMDM experiments. After 4 h, media was removed and cells were lysed with 40 µl TNT lysis buffer (20 mM Tris, pH 8, 200 mM NaCl, 1% triton-100). Lysates were then injected with 40 µl firefly luciferin substrate (Biosynth) and luminescence was measured using a SpectraMax L plate reader (Molecular Devices).

### qPCR

For qPCR experiments using mouse tissue, 50 µl of organ homogenates was added to 600 µl of RLT buffer (Qiagen) containing 1% β-mercaptoethanol and frozen at −80°C. Frozen lysates were later thawed and RNA was purified and treated with DNAse, according to the manufacturer’s protocol using an RNeasy kit (Qiagen). RNA abundance was quantified using a NanoDrop ND-1000 and 200 ng RNA was *in vitro* transcribed (ProtoScript II, NEB, M0368S) and diluted 10x in sterile nuclease-free water. Real-time PCR was performed using SYBR Green (Thermo, A25742), 2 µl of cDNA, and 1 µM each of the following oligonucleotides: actin F: GGCTGTATTCCCCTCCATCG; actin R: GTCACCCACATAGGAGTCCTTC; IFN-β F: AGCTCCAAGAAAGGACGAACAT; and IFN-β R: CCCTGTAGGTGAGGTTGATCTT. For normalization, values for IFN-β were divided by values for actin. Measurements were acquired with a QuantStudio 5 real-time qPCR machine (Applied Biosystems).

## QUANTIFICATION AND STATISTICAL ANALYSIS

Statistical parameters and significance are reported in the figures and the figure legends. Data are determined to be statistically significant when p<0.05 by an unpaired two-tailed Student’s T-Test an ANOVA with multiple comparisons post-hoc test. For *in vivo* PFU data, data are determined to be statistically significant when p<0.05 by a Mann-Whitney *U* test. Asterisks denote statistical significance as: *, p<0.05; **, p<0.01; ***, p<0.001; ****, p<0.0001, compared to indicated controls. For animal experiments, bars denote medians. Error bars indicate standard error (SE). All other graphical representations are described in the figure legends. Statistical analyses were performed using GraphPad PRISM.

---

**Table S1. Related to Figure 3. RNA-seq analysis of genes regulated by IRFs during R. parkeri infection and IFN-I treatment.**

