## Supplemental Figures for "Inflammasome-mediated antagonism of type I interferon enhances *Rickettsia* pathogenesis"

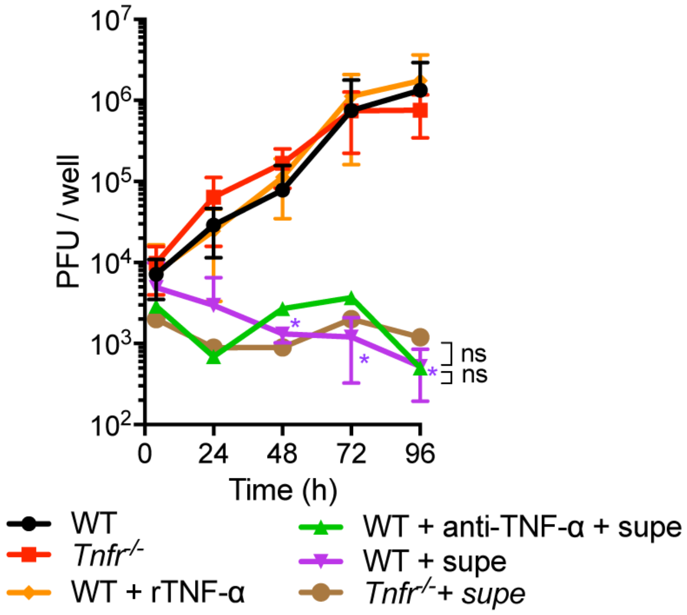


**Figure S1. Related to Figure 1: TNF-α does not affect *R. parkeri* growth in BMDMs**

Measurement of *R. parkeri* abundance in BMDMs. Cells were infected at an MOI of 0.2 and PFUs were measured over time. Statistical comparisons were made between each sample and WT B6 cells at each time point. Data are the combination of at least three separate experiments and are expressed as means + SEM. “Supe” indicates 200 ul of conditioned supernatant collected at 24 hpi from *Casp1$^{-/-}$Casp11$^{-/-}$* BMDMs infected at an MOI of 1. All statistical analyses were performed using a Student’s T-test. *p<0.05. ns = not significant.

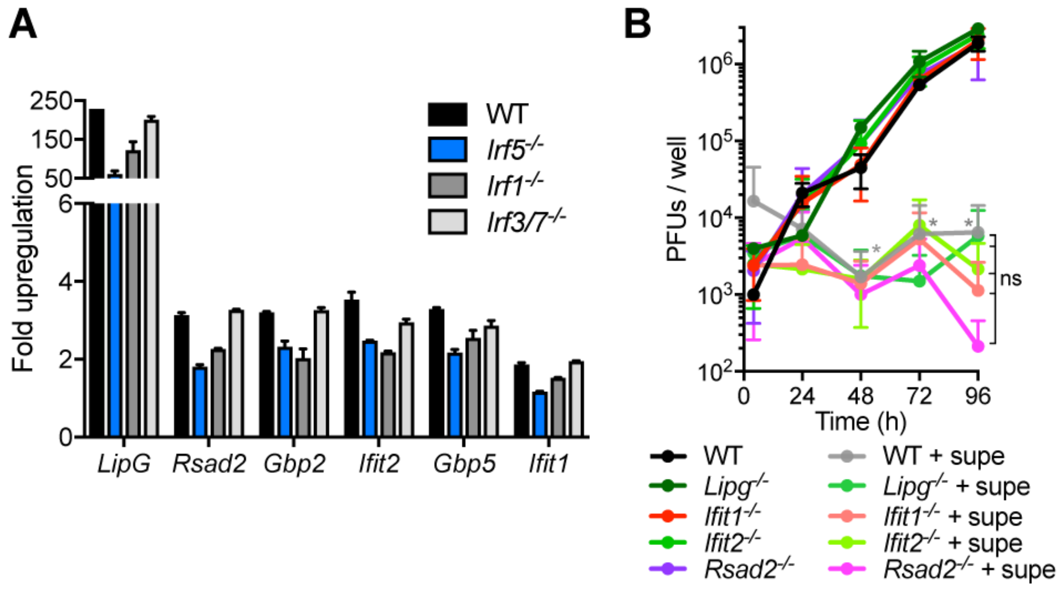


**Figure S2. Related to Figure 3. Candidate anti-rickettsial genes regulated by IRF5, as analyzed by qPCR and PFU growth curve**

**A)** qPCR of the indicated genes identified from RNA-seq. Cells were infected with *R. parkeri* at an MOI of 2.3, as determined by immunofluorescence microscopy, and treated with 10,000 U rIFN-β (PBL) at T=0. RNA was harvested at 12 hpi, reverse transcribed into cDNA, and analyzed with qPCR. To determine fold upregulation, each sample was compared to infected WT cells that were not treated with IFN-β. The abundance of each gene product was normalized to actin. Data are the average of two separate experiments. **B**) *R. parkeri* growth in BMDMs. BMDMs were infected with *R. parkeri* at an MOI of 0.2 and were PFUs measured over time. Data are at least three combined experiments and are expressed as means $\underline{+}$ SEM. “Supe” indicates 500 ul of supernatant collected at 24 hpi from *Casp1*$^{-/-}$*Casp11*$^{-/-}$ BMDMs infected at an MOI of 1. Statistical analyses were performed using a Student’s T-test. *p<0.05. ns = not significant.

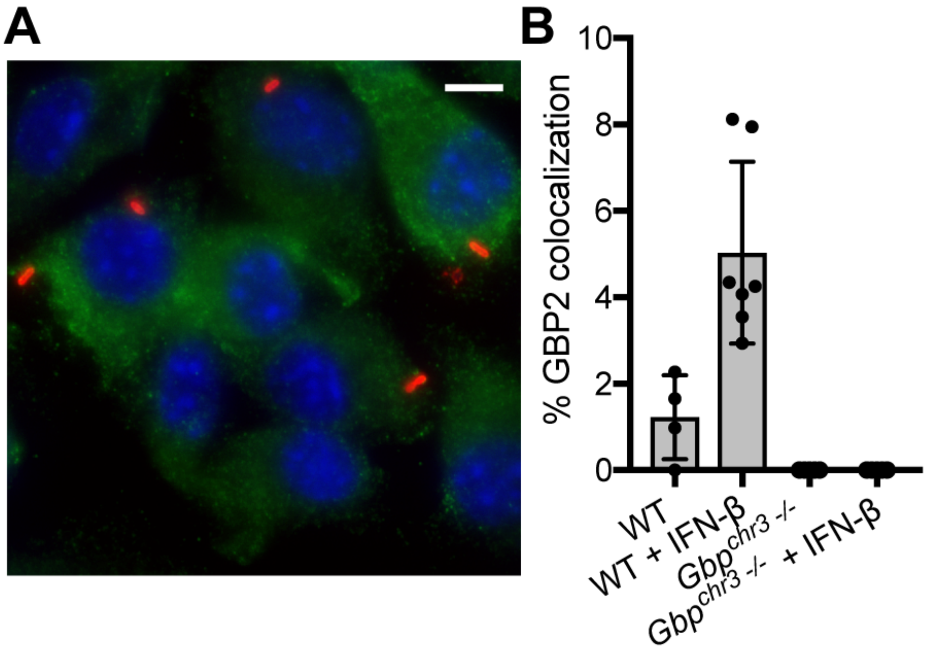


**Figure S3. Related to Figure 4. No colocalization is observed between GBP2 and *R. parkeri* in *Gbp*$^{chr3-/-}$ BMDMs**

**A**) A representative image of *R. parkeri* in *Gbp*$^{chr3-/-}$ BMDMs, as stained with an anti-GBP2 antibody. Immunofluorescence microscopy was used to evaluate GBP2 localization. Green staining is α-GBP2; red staining is α-*Rickettsia*; blue staining is DAPI. The scale bar is 5.6 µm. **B**) Quantification of bacteria that co-localized with GBP2. Each data point represents an individual experiment, and each experiment consists of at least 10 separate images, and each image contained approximately 20 bacteria. At least 3 separate experiments were performed in *Gbp*$^{chr3-/-}$ BMDMs. No bacteria were observed in *Gbp*$^{chr3-/-}$ BMDMs that localized to GBP2 staining.

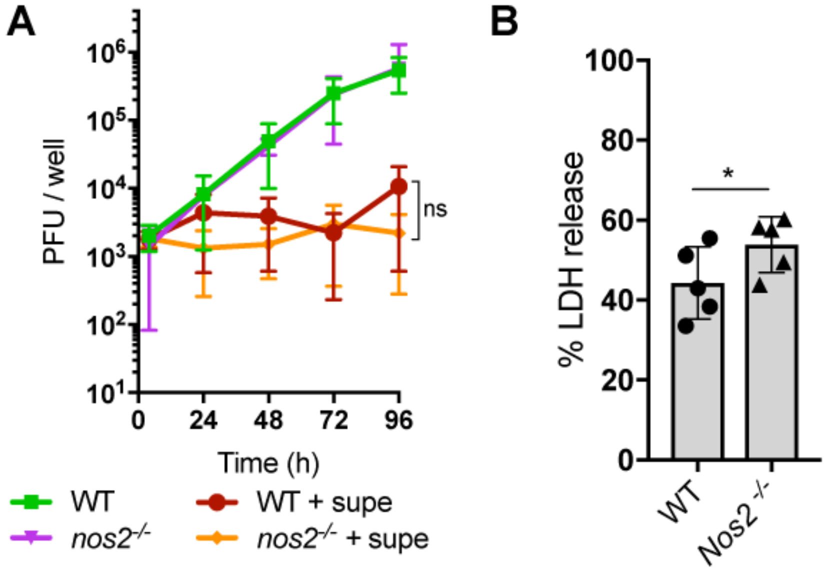


**Figure S4. Related to Figure 4. *R. parkeri* do not grow better upon IFN-I treatment in *Nos2*$^{-/-}$ BMDMs, however these cells undergo higher cell death than WT cells**

**A**) *R. parkeri* growth in BMDMs. The indicated BMDMs were infected with *R. parkeri* at an MOI of 0.2 and PFUs measured over time. Data represent at least three separate experiments and are expressed as means $\pm$ SEM. "Supe" indicates 200 ul of supernatant collected at 24 hpi from *Casp1*$^{-/-}$*Casp11*$^{-/-}$ BMDMs infected at an MOI of 1. **B**) Host cell death upon infection with *R. parkeri*. Supernatants were collected from the indicated cells that were infected at an initial MOI of 1 and the amount of LDH was measured at 24 hpi. Statistical analyses were performed using a Student's T-test. *$p<0.05$. ns = not significant.

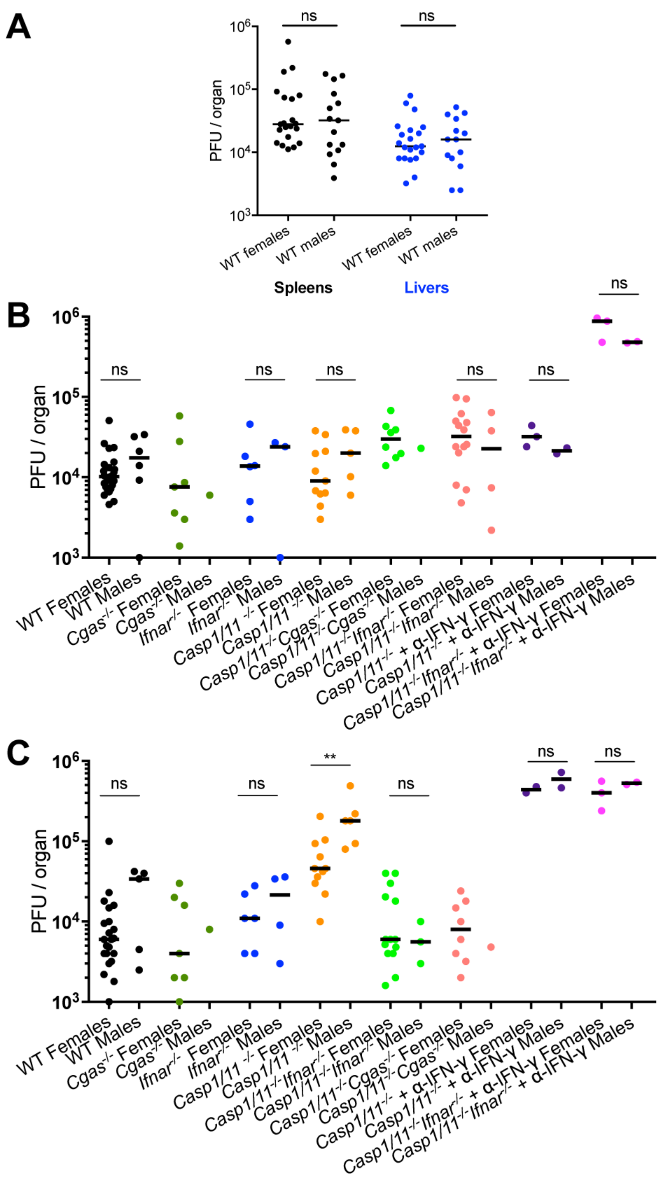


**Figure S5. Related to Figure 5. *R. parkeri* growth in male and female mice.**

**A)** *R. parkeri* burdens in C57Bl/6 male and female mice at 48 hpi. Mice were infected i.v. with $10^7$ *R. parkeri* and bacterial burdens were evaluated using PFU assays in the indicated organs at 48 hpi. **B,C**) *R. parkeri* burdens in C57Bl/6 male and female mice at 72 hpi. Each mouse was infected with $10^7$ *R. parkeri* i.v. and bacterial burdens were evaluated in spleens and livers at 72 hpi. The combined data are shown in Figure 5. Statistical analyses for in vivo experiments were performed using a Mann-Whitney U test. Bars denote medians. **p<0.01 and ns=not significant.
